# Unraveling Subcellular Ultrastructure with Cyclically Multiplexed Expansion Microscopy

**DOI:** 10.64898/2026.01.01.697161

**Authors:** Seweryn Gałecki, Bo-Jui Chang, Felix Zhou, Qionghua Shen, Daniel Stoddard, Bingying Chen, Daniela Nicastro, Reto Fiolka, Kevin M. Dean

**Affiliations:** Lyda Hill Department of Bioinformatics, UT Southwestern Medical Center, Dallas, TX 75390, USA; Cecil H. and Ida Green Center for Systems Biology, UT Southwestern Medical Center, Dallas, TX 75390, USA; Department of Systems Biology and Engineering, Silesian University of Technology, Akademicka 16, 44-100 Gliwice, Poland; Department of Biophysics, UT Southwestern Medical Center, Dallas, TX 75390, USA; Department of Cell Biology, UT Southwestern Medical Center, Dallas, TX 75390, USA

## Abstract

Despite advances in fluorescence microscopy, spectral overlap and limited resolution hinder the dense mapping of the cellular ultrastructure. To overcome these challenges, we developed Cy-ExM, a high-plex imaging strategy that integrates optimized cryo-fixation for antigen preservation, expansion microscopy, and iterative immunofluorescence labeling. Using oblique plane microscopy, we perform three-dimensional super-resolution imaging of 20 biological targets encompassing the full cellular volume of individual mammalian cells.

## Main Text

The spatial organization of proteins and organelles within eukaryotic cells governs a vast amount of fundamental biological processes such as replication, metabolism, and intra– and extracellular signaling. Fluorescence microscopy is a widely applicable method for visualizing subcellular structures because it provides high molecular specificity and relatively rapid, multi-target imaging, whereas electron microscopy excels at revealing ultrastructural and cellular context but is generally lower throughput and, in conventional implementations, depends on broad-spectrum contrast mechanisms such as heavy metal staining^1^. Nonetheless, spectral overlap and the limited availability of orthogonal antibody pairs constrain the number of molecular targets that can be simultaneously visualized in fluorescence microscopy. While spectral imaging and unmixing techniques can computationally separate overlapping fluorophores, their accuracy diminishes at low photon counts where noise is non-negligible. As a result, modern multiplexed imaging approaches, such as t-CyCIF^2^, IBEX^3^, and CODEX^4^, have adopted cyclic labeling strategies to dramatically expand the number of visualized targets. Together, these methods have provided critical insights into physiological and pathological states, particularly in tissue contexts^5^.

However, understanding cellular function requires imaging molecular structures at cellular and sub-cellular scales, where key biological processes such as protein complex assembly, cytoskeletal organization, and organelle interactions occur. Super-resolution light microscopy has greatly advanced our ability to visualize biological structures at sub-diffraction-scales^6^, but practical limitations hinder its application in multiplexed imaging. Techniques like localization microscopy and stimulated emission depletion rely on specialized fluorophores, and their inherently slow acquisition speeds constrain imaging to shallow 3D volumes^7^. Even with state-of-the-art DNA-PAINT-based multiplexing methods, imaging is typically limited to about a dozen targets and shallow (∼1 micron) imaging depths^8^. Overcoming these challenges requires an integrated imaging strategy that combines volumetric super-resolution with high-plex molecular detection in thick specimens, enabling a more comprehensive understanding of cellular organization.

One promising strategy for multiplexed, nanoscale imaging involves embedding specimens in a hydrogel scaffold that simultaneously stabilizes the sample mechanically and renders it optically transparent. However, embedding and polymerization steps, often coupled with proteolytic softening to relax tissue mechanics, can mask or remove antigens, reducing sensitivity across a range of molecular targets^9^. Immunostaining can be performed either before or after hydrogel embedding, but each approach requires a distinct workflow and involves trade-offs in labeling efficiency, signal retention, and multiplexing capacity (See **Supplementary Note 1**). Despite these challenges, techniques such as CLARITY^10^, SWITCH^11^, and Expansion Microscopy^12^, have been adapted for cyclic labeling strategies. To date, however, most cyclic ExM-style applications have emphasized synaptic marker panels and synapse classification^12^, in part because extending these workflows to comprehensive, cell-biological mapping of diverse organelles and protein assemblies requires substantially higher preservation of ultrastructure and epitope integrity across repeated processing steps. In practice, cumulative losses in antigenicity, structural distortions introduced during mechanical homogenization, and signal dilution in expanded gels have limited broad application to multi-organelle nanoscale mapping. Consequently, a cell-biologically focused approach that fully integrates cyclic multiplexing with expansion microscopy while preserving ultrastructure across cycles has yet to be demonstrated.

To overcome these limitations, we introduce Cyclically Multiplexed Expansion Microscopy (Cy-ExM), a strategy for high-sensitivity, nanoscale, three-dimensional visualization of subcellular architectures. Our method combines cryofixation by guillotine-based plunge-freezing, which minimizes specimen damage (**Figure S1**), with acetone-based freeze substitution, protein entanglement, and heat-induced denaturation to relax intermolecular interactions and mechanically homogenize the specimen for isotropic expansion^13^ (**Figure 1a-b, Supplementary Note 2**). Although denaturation disrupts native protein conformation, it maintains the relative spatial arrangement of proteins and minimizes antigen loss compared with conventional chemical fixation and expansion workflow^14^ (**Figure S2**). Leveraging this approach, specimens routinely achieved 4.2-fold linear expansion and were compatible with volumetric imaging on both conventional fluorescence microscopes (**Figure 1c)** and an inverted oblique plane microscope (OPM) optimized for large, aqueous samples, with a large field of view and a high numerical aperture (NA 1.1) water-dipping objective^15^. Combined, our sample preparation, labeling, and OPM achieved ∼95 nm lateral and ∼290 nm axial resolution prior to deconvolution, and ∼70 nm lateral by ∼220 nm axial resolution after deconvolution, respectively, while enabling iterative multiplexed mapping of up to 20 targets. Relative to standard immunofluorescence, Cy-ExM revealed significantly greater structural detail, including the visualization of mitochondrial cristae and continuous, well-resolved microtubule filaments throughout the cytoskeletal network in mouse embryonic fibroblast (MEF) cells (**Figure 1d-g, Figure S3**).

**Figure 1.**
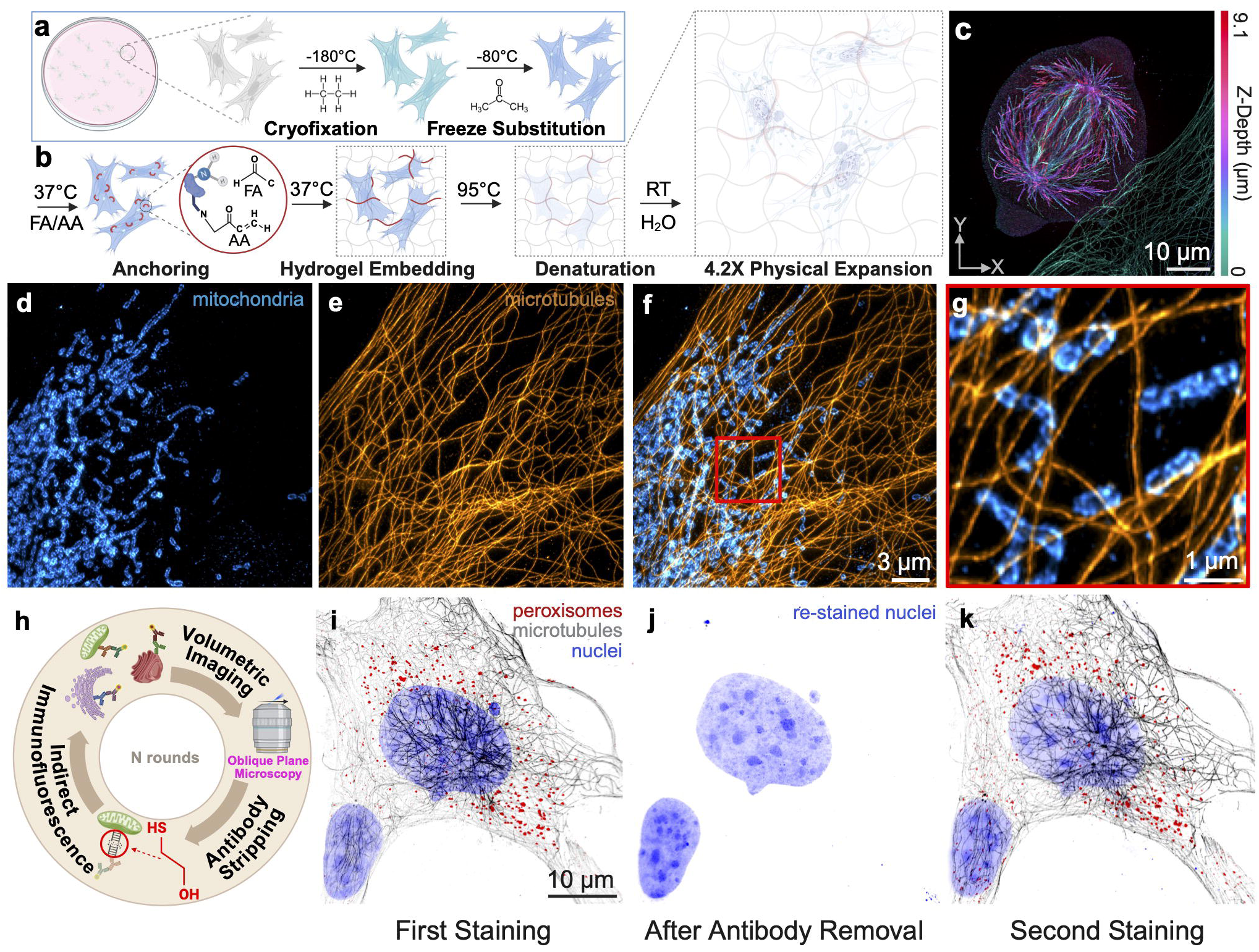
Cyclically Multiplexed Expansion Microscopy Workflow. **(a-b)** Overview of sample preparation for Cy-ExM. **(a)** Cultured cells were rapidly plunge-frozen in liquid ethane at –180°C and freeze-substituted into acetone at –80 °C to preserve ultrastructure and antigenicity. **(b)** After rehydration, cells were chemically anchored with formaldehyde and acrylamide, embedded in a swellable hydrogel, denatured at 95 °C to expose epitopes, and isotropically expanded in deionized water. **(c)** Depth-encoded maximum-intensity projection of a cryo-fixed, 4.2× expanded MEF undergoing mitosis, in which color indicates the axial (Z) position. **(d–g)** Representative images of cultured MEFs processed using a single round of the Cy-ExM workflow. **(d)** Mitochondria channel only (cyan). **(e)** Microtubules channel only (orange). **(f)** Merged view of microtubules (orange) and mitochondria (cyan). **(g)** Magnified inset corresponding to the boxed region in **(f)**. **(h)** The full Cy-ExM workflow: **v**isualization of cellular ultrastructure was performed by iterative post-expansion indirect immunofluorescence, volumetric imaging with an oblique plane or spinning disk confocal microscope, and antibody stripping using β-mercaptoethanol. **(i–k)** Example of iterative imaging across rounds. Cells were first labeled for peroxisomes, microtubules, and nuclei and imaged; antibodies were then stripped, nuclei were re-stained and re-imaged as a reference; cells were subsequently re-labeled for peroxisomes, microtubules, and nuclei and imaged again. Deconvolved maximum-intensity projections are shown. All scale bars are corrected for the expansion factor.

Importantly, this protocol enabled repeated cycles of labeling and volumetric imaging of the same specimen (**Figure 1h**) while preserving subcellular architecture and maintaining a reproducible 4.2× linear expansion coefficient across successive rounds. We used nuclear DNA staining as a robust fiduciary marker to quantify expansion repeatability and confirm preservation of fine nuclear features (e.g., nucleoli) across cycles (**Figure S4**). To further evaluate the fidelity of iterative labeling, cryo-fixed MEFs were embedded in a swellable hydrogel, denaturated, and partially expanded (∼2×) for immunolabeling of peroxisomes, DNA, and microtubules, and then fully expanded (4.2×) for volumetric imaging (**Figure 1i-k, Movie S1, Tables S1 and S2**). After the first round of imaging (**Figure 1i**), antibodies were gently stripped using β-mercaptoethanol to reduce disulfide bonds and facilitate antibody removal^12^; complete loss of antibody signal in the stripped state confirmed effective stripping (**Figure 1j**). Cells were then re-labeled with the same antibody set and re-imaged (**Figure 1k**).

Although visual inspection showed strong agreement across labeling rounds, we next quantified how faithfully subcellular features aligned in 3D by quantitatively evaluating registration accuracy between rounds using the same marker sets. Registration was performed using the microtubule channel, which is distributed throughout the cellular interior and is sufficiently dense to robustly constrain the registration transform. To ensure accurate alignment between rounds, we applied a pyramidal registration routine that increased in complexity over successive steps: beginning with simple translation, followed by rigid body, similarity, and ultimately two rounds of affine registration (**Figure S5a; Methods**). Registration accuracy in 3D was evaluated using quantitative metrics, including normalized cross-correlation, mutual information, and structural similarity index (**Supplementary Note 3**). Decomposition of the affine matrix showed that round-to-round transformations were small, with near-zero rotation and shear, and only slight anisotropic scaling (**Table S3**). While affine alignment achieved strong global consistency, quantitative similarity metrics showed little additional improvement following non-linear warping (**Table S4**). Nevertheless, inspection of the registered volumes revealed subtle but spatially localized misalignments that were not captured by global intensity and structure-based metrics. These residual local distortions were effectively corrected by applying a modest non-linear warp (**Figure S5b-c, Movie S2**), suggesting that non-linear registration primarily refines local geometric consistency rather than improving global similarity scores, and resulting in high-fidelity alignment between individual imaging rounds.

To determine whether this level of accuracy was maintained across the entire cyclic workflow, not just between two rounds, we next evaluated the consistency of registration and labeling across five Cy-ExM cycles. Image registration was performed using the microtubule channel, and registration accuracy was independently assessed in the mitochondria channel, which was not used during alignment, by detecting mitochondrial puncta and computing mutual nearest-neighbor–paired distances as a target registration error (TRE)-like proxy (**Figure S6**). Using this approach, we observed a root mean square displacement of ∼124 nm across cycles (**Table S5**). Nearest-neighbor pairing provides a straightforward estimate of residual misalignment, but for spatially heterogeneous targets such as mitochondria, incorrect correspondences can occur when nearby puncta are ambiguously matched. To obtain a more conservative, globally consistent estimate, we additionally matched mitochondrial features using descriptor-based correspondences and robustly fit a global similarity transform to the resulting pairs, estimating the TRE proxy from inlier matches. This approach yielded a mean cross-validated TRE proxy of 91 and 100 nm, after linear and nonlinear registration, respectively. Together, these analyses indicate that Cy-ExM preserves high spatial fidelity across iterative cycles, with residual inter-round misalignment on the order of ∼100 nm throughout the entire field of view.

We next quantified round-to-round staining variability with a spinning disk confocal microscope using a 20× NA 0.75 air objective (**Figure S7**). Peroxisome signal was quantified by measuring the background-corrected mean intensity within regions of interest for seven cells; after stripping, the peroxisome intensities dropped to near-background levels (i.e., comparable to regions outside the cell), yielding signal-to-noise values close to unity. Notably, peroxisome labeling intensity did not monotonically decrease across cycles; instead, it increased from rounds 1–4 and then modestly decreased in round 5 while remaining higher than rounds 1–3, consistent with cycle-dependent changes in staining efficiency rather than cumulative antigen loss. Consistent with this interpretation, a cycle-permutation experiment showed that antigenicity was independent of labeling order (**Figure S8**), indicating that epitopes remain accessible even when targets are stained in late rounds. Because antibodies are gently removed using redox chemistry and the specimen is mechanically reinforced by the hydrogel, the apparent labeling quality was not sensitive to staining order, as judged by consistent staining patterns and low background across cycles. Consequently, we hypothesized that Cy-ExM should tolerate cycle counts comparable to established cyclic immunofluorescence workflows (e.g., 10-20 cycles, depending on tissue resilience^2,3,16^). To evaluate this, we simulated a high-cycle experiment by subjecting a gel to ten consecutive rounds of stripping, consisting of 45 minutes in β-mercaptoethanol at 95°C, with staining and imaging performed at the beginning, middle, and end of the workflow. Cells were labeled for actin filaments, microtubules, mitochondrial matrix, endoplasmic reticulum (ER), and DNA, and all structures remained well preserved even after ten rounds of stripping (**Figure S9**).

To demonstrate the feasibility and practical utility of Cy-ExM, we next applied the method to visualize 20 distinct molecular targets within the same MEF cell across eleven iterative imaging cycles (**Figure 2a-b**, **Movie S3, Table S6**). This multiplexed dataset captured the major cellular organelles, including the ER, Golgi apparatus, mitochondria, lysosomes, and peroxisomes, as well as nuclear components such as the nuclear envelope, nuclear pore complexes, and fibrillarin). In addition to membrane-bound organelles, cytoskeletal networks comprising actin filaments, vimentin, and microtubules, were robustly resolved, reflecting the broad epitope compatibility enabled by cryo-fixation and post-expansion labeling. Analysis of the full dataset confirmed that geometric stability was maintained across labeling cycles at scale: decomposition of affine transforms revealed highly consistent round-to round alignment with minimal variability in scale and shear, and non-linear warping remained modest, indicating limited cumulative distortion (**Table S7**). Given the multiplexing demonstrated here, Cy-ExM enables direct comparison of multiple cellular structures within the same cell, avoiding reliance on population-averaged measurements across separate specimens^17^. To illustrate this capability, we organized datasets acquired from distinct cells into functionally coherent views of subcellular architecture (**Figure 2c–f, Movie S3 and S4**), highlighting distinct regions of interest in which targets were grouped by primary biological role, including the secretory pathway, cytoskeletal network, metabolic organelle network, and nuclear architecture. This integrative representation preserves the full molecular context of each structure while maintaining sufficient spatial resolution to resolve nanoscale features, such as individual NUP p62 nuclear pore complexes within the nuclear envelope (**Figure 2f, Figure S10**). Together, these capabilities position Cy-ExM as a practical route toward constructing high-content nanoscale atlases of cellular ultrastructure, enabling systematic mapping and quantitative analysis of organelle organization and interactions within a single, intact cell.

**Figure 2.**
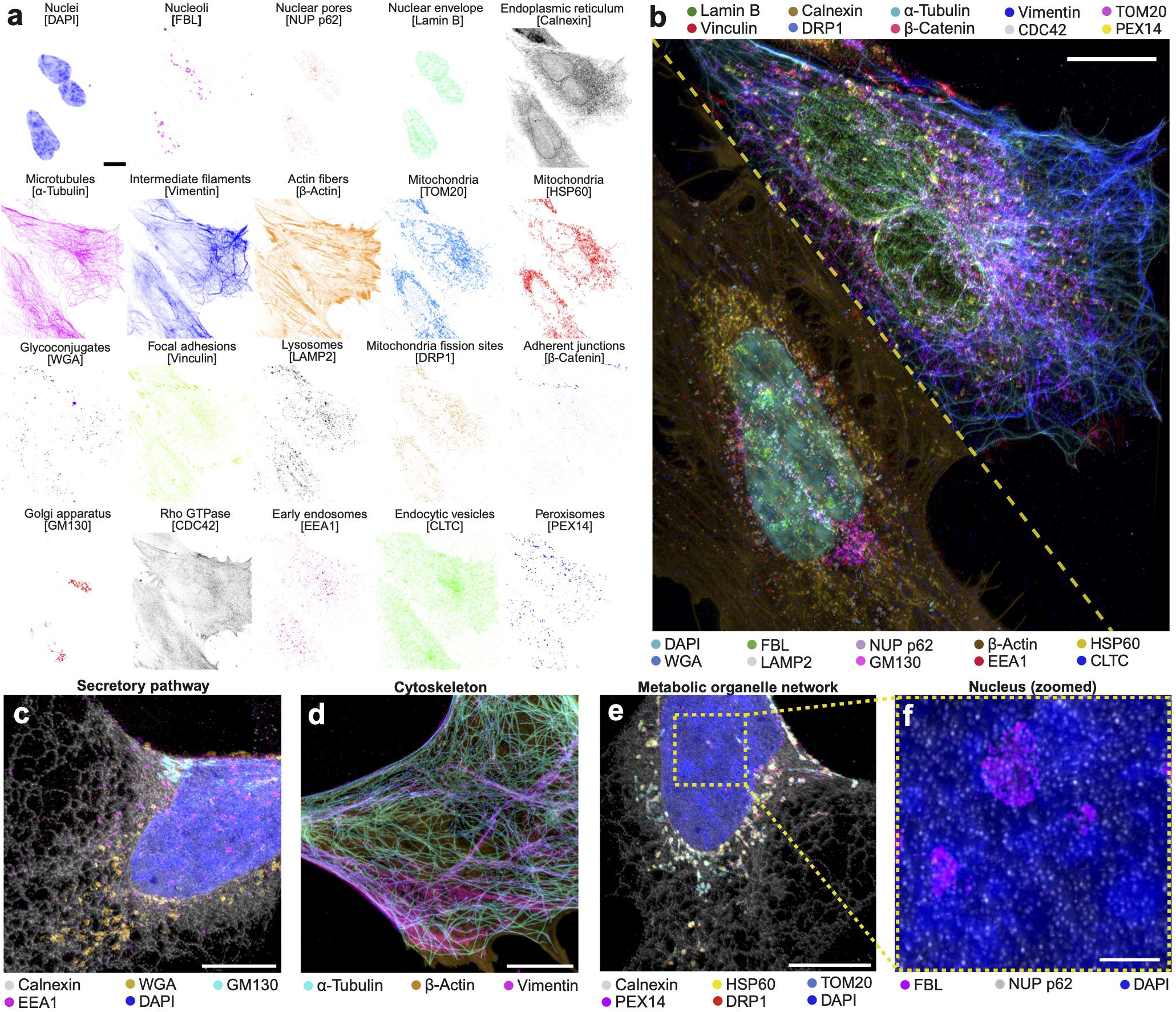
Twenty-plex volumetric Cy-ExM imaging of subcellular organization. **(a)** Maximum-intensity projections (MIPs) of all 20 cellular targets imaged in the same cell using the Cy-ExM workflow; individual channels are displayed with inverted lookup tables to enhance visibility of fine structural features (target labels indicated above each panel). Inter-round registration was performed using the microtubule channel as the structural reference. **(b)** Corresponding merged MIPs of the same cell are shown as two composite subpanels, each displaying 10 targets (lookup tables indicated by colored dots). **(c–f)** Data from a different cell, with targets grouped into four functional classes: **(c)** secretory pathway, **(d)** cytoskeletal components, **(e)** metabolic organelle network, and **(f)** nuclear organization. All images were acquired volumetrically using an oblique plane microscope and deconvolved prior to visualization. Scale bars: **(a)** 10 µm; **(b)** 20 µm; **(c–e)** 5 µm; **(f)** 2 µm (all corrected for the expansion factor).

One compelling application of Cy-ExM is to quantify coordinated spatial organization of proteins and organelles within single cells. To test this, we compared the radial distributions of multiplexed markers from the nuclear boundary to the cell periphery by computing per-marker intensity profiles normalized by cell size (**Figure 3a–b; Methods**). Because distances are normalized, these profiles provide a rotationally invariant, shape-independent representation of spatial organization that enables direct comparisons across cells. Markers associated with the same organelles exhibited similar radial profiles, and clustering of these profiles recovered expected groupings of mitochondrial, nuclear, and cytoskeletal components (**Figure 3c; Figure S10**). These relationships were reproducible across independent cells (**Figure 3d**), demonstrating that Cy-ExM supports quantitative, single-cell comparisons of subcellular organization and provides a practical framework for testing how organelle organization changes across cell states and perturbations.

**Figure 3.**
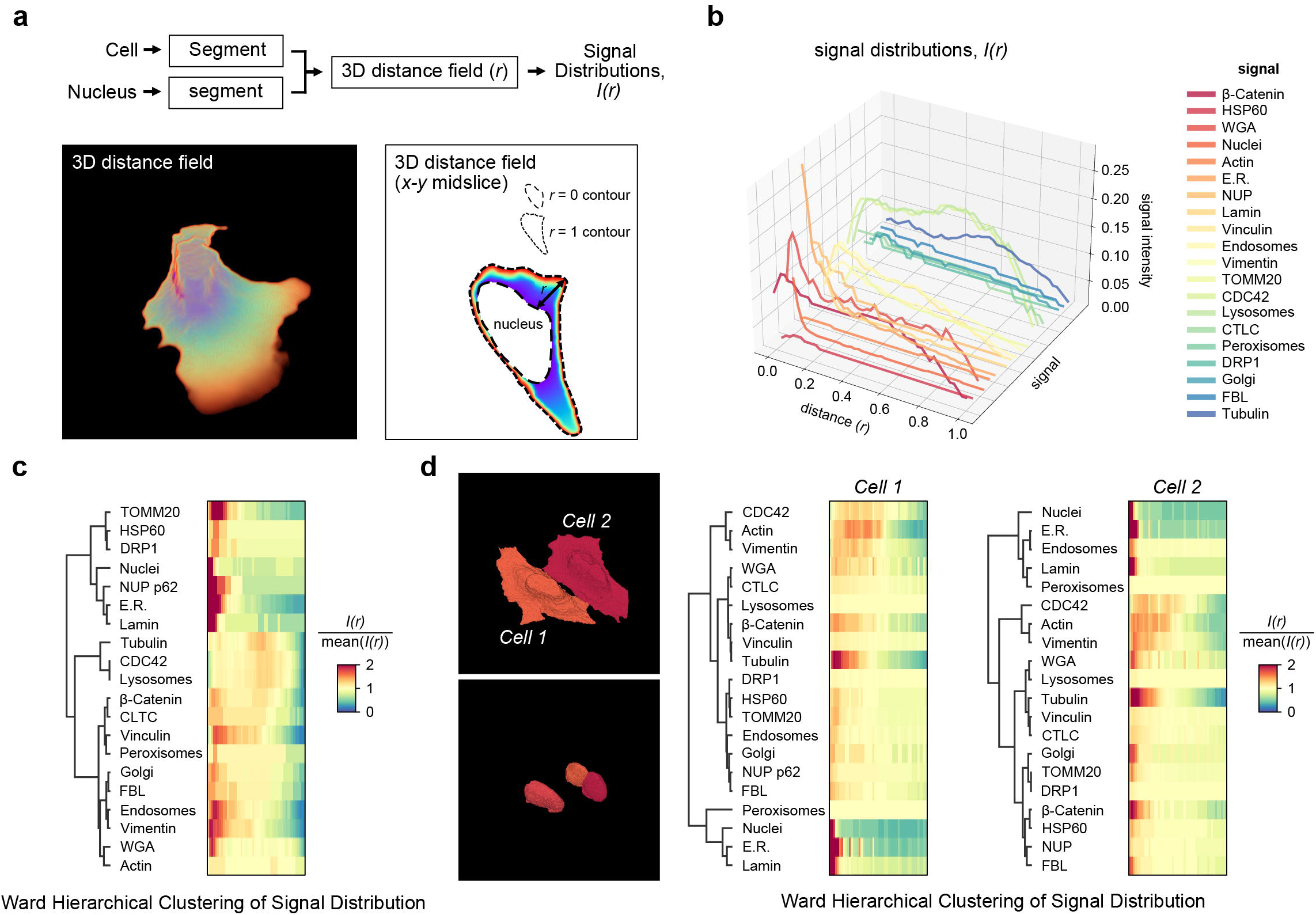
Quantifying co-organization of subcellular architectures using normalized radial signal distributions. **(a)** Analysis workflow: the cell and nuclear volumes are segmented to compute a 3D normalized distance field (*r*), where *r* = 0 corresponds to the nuclear boundary and *r* = 1 to the cell periphery, enabling calculation of a rotationally invariant, shape-normalized signal distribution, *I(r)*. Representative renderings of the 3D distance field and an *x–y* midslice are shown, with example *r* = 0 and *r* = 1 contours indicated. **(b)** Normalized signal distributions, *I(r)*, for 20 markers in the same cell shown in Figure 2c–f. **(c)** Ward hierarchical clustering of the radial signal distributions from the single-cell 20-plex dataset, displayed as a dendrogram and heatmap of *I(r)/mean(I(r))*. **(d)** Two additional segmented cells (shown in Figure 2a–b) and the corresponding Ward hierarchical clustering of their radial signal distributions (Cell 1 and Cell 2), demonstrating reproducible organization patterns across cells.

Taken together, these results establish Cy-ExM as a high-fidelity platform for high-cycle, three-dimensional multiplexed imaging that preserves ultrastructural context, antigenicity, and spatial fidelity across repeated labeling rounds (**See Supplementary Note 4, Limitations and Future Directions**). By integrating cryo-preserved sample preparation with expansion microscopy and iterative immunolabeling, Cy-ExM enables quantitative, whole-cell volumetric super-resolution maps of multi-protein organization using standard fluorescence microscopes. We anticipate that this combination of ultrastructural preservation and multiplexed nanoscale readouts will make it possible to construct systematic atlases of subcellular architecture and to measure how organelle organization and compartmental co-organization evolve in increasingly complex specimen contexts.

## Acknowledgements

K.M.D. is supported by the Cancer Prevention and Research Institute of Texas RP250571, the NCI U54CA268072, and the NIGMS RM1GM145399. R.F. is supported by NCI U54CA268072, NIGMS R35GM133522, and NIBIB R01EB035538, D.N. is supported by NIGMS R01GM083122 and R01GM154131. We acknowledge Dr. Tadamoto Isogai and Dr. Mike Henne for providing a portion of the primary antibodies for validation and staining of cellular structures, as well as Dr. Dana Reed for administrative support. We would also like to acknowledge the Quantitative Light Microscopy Core, which is supported by the Harold C. Simmons Cancer Center at UT Southwestern Medical Center (NCI 1P30CA142543) and the NIH Shared Instrumentation Awards (NIGMS S10OD028630). We thank the University of Texas Southwestern Medical Center Cryo-EM Facility, which is partially funded by the Cancer Prevention and Research Institute of Texas Core Facility Support Award RP220582. Figures in this manuscript were created with BioRender.

## Author Contributions

S.G.: Methodology, Validation, Formal Analysis, Investigation, Writing (original draft, review, and editing), Visualization. B.-J.C.: Methodology, Investigation, Writing (review and editing). Q.S.: Methodology, Validation, Investigation, Writing (review and editing). D.S.: Methodology, Writing (review and editing). B.C.: Methodology, Investigation, Writing (review and editing). D.N.: Methodology, Writing (review and editing). F.Z.: Methodology, Writing, Investigation. R.F.: Resources, Investigation, Writing (review and editing). K.M.D.: Conceptualization, Methodology, Software, Formal Analysis, Writing (original draft, review, and editing), Resources, Visualization, Supervision, Project Administration, Funding Acquisition.

## Declaration of generative AI and AI-assisted technologies in the manuscript preparation process

During the preparation of this work the authors used ChatGPT in to improve grammar and readability of the text. After using this service, the authors reviewed and edited the content as needed and takes full responsibility for the content of the published article.

## ONLINE METHODS

### Cell culture

Mouse embryonic fibroblasts (MEFs) cells were cultured under standard conditions in Dulbecco’s Modified Eagle’s Medium (DMEM, Gibco) supplemented with 10% fetal bovine serum (FBS, Gibco), 100 U/mL penicillin, and 100 µg/mL streptomycin. The cells were maintained at 37°C in a humidified incubator with 5% CO and cultured in a 12-well plate on 5-mm glass coverslips pre-rinsed with 70% ethanol.

### Chemical fixation

To compare fixation methods (**Figure S2**), MEF cells were rinsed with 1× Phosphate-Buffered Saline (PBS) and incubated in: (a) 4% paraformaldehyde (PFA) at RT for 15 min, (b) 4% PFA supplemented with 0.2% Glutaraldehyde (GA) at RT for 15 min, (c) pre-heated 37°C PEM buffer [80 mM PIPES, 5 mM EGTA, 2 mM MgCl_2_, (pH: 6.8)], supplemented with 0.3% Triton-X and 0.125% GA for 30 s, fixed in a PEM^18^ buffer supplemented with 2% PFA for 15 min at 37°C and washed three times with 1× PBS, 2 min each, or (d) −20°C chilled methanol for 5 min. In all other cases, the cells were subjected to cryofixation.

### Cryofixation

Cells were cryo-fixed by plunge freezing followed by solvent fixation during freeze substitution using an established, slightly modified protocol^14^. Briefly, a glass coverslip with adherent cells was held vertically with forceps and gently blotted with cotton-tipped applicators. Using a homemade pneumatic plunge freezing device, the coverslip was then rapidly plunged into liquid ethane that was chilled in a liquid N_2_-filled Vitrobot dewar (Thermofisher). Next, the coverslip was quickly immersed in liquid nitrogen and transferred into an Eppendorf tube pre-cooled in liquid nitrogen and containing 1.5 mL of frozen acetone. The tubes were stored in liquid nitrogen until further processing. For freeze substitution, the tubes with coverslips were left overnight in a metal block that was cooled with dry ice and liquid nitrogen in a cold room, allowing gradual warming to −80°C and thawing of the acetone. Next day, the dry ice was removed, and the tubes were incubated in the metal block with lids open at room temperature for ∼1 h until the acetone reached 0°C, completing solvent fixation. Samples were then rehydrated through successive ethanol:water baths in the following gradients: 100% (x2), 95% (x2), 70%, 50%, and 25% EtOH, and finally 1× PBS, for 5 min each. Rehydrated samples were stored at 4°C in 1× PBS with 0.02% (w/v) sodium azide (NaN_3_) to prevent microbial and fungal growth.

### Ultrastructure Expansion Microscopy

Cryo-fixed cells were rinsed three times with 1× PBS to remove NaN_3_ and underwent anchoring in 1× PBS containing 1.4% formaldehyde (FA, Sigma-Aldrich) and 2% acrylamide (AA, Bio-Rad) for 3 h at 37°C without agitation. Cells were then incubated in a monomer solution [23% sodium acrylate (SA, AmBeed), 10% AA, 0.1% N,N′-methylenbisacrylamide (BIS, Sigma-Aldrich), 5% Tetramethylethylenediamine (TEMED, Bio-Rad), 1× PBS and Mili-Q water] for 30 min on ice to ensure uniform monomer infusion of the cells. After monomer infusion, the glass coverslips were placed cell-side down onto an ice-cooled (4°C) gelation chamber containing ∼15 μL monomer solution with 0.5% ammonium persulfate (APS, Bio-Rad) for 10 min. After initial polymerization on ice, the samples were transferred to 37°C and incubated for 45 min under humidified conditions for complete gel formation. To relax native protein assemblies and enable uniform expansion, hydrogels were next transferred to a vial containing denaturation buffer [200mM SDS, 200mM NaCl, 50mM Tris-Base, and Mili-Q water, (pH 9)], and incubated for 1.5 h in preheated 95°C oven or thermoblock. At this stage, gels expanded ∼twofold and detached spontaneously from the coverslips. Finally, hydrogels were immersed in Mili-Q water for at least 1 h, with water exchanged every 20 min or left overnight at RT to ensure complete physical expansion and removal of denaturation agents. The size of the hydrogel was measured to calculate the macro-scale expansion factor, after which it was transferred to 1× PBS to partially shrink prior to labeling. For storage, hydrogels were cut into smaller pieces and stored at 4 °C in 1× PBS containing 0.02% (w/v) NaN to prevent microbial and fungal growth.

### Highly Multiplexed Labeling of Expanded Cells

All incubations were performed at room temperature with constant agitation unless otherwise specified. To enhance epitope accessibility and improve signal-to-noise, hydrogels shown in **Figure 2** and **Supplementary Figure 10** were pre-treated in denaturation buffer supplemented with 100 mM β-mercaptoethanol (BME) and incubated at 95 °C for ∼30 min. Prior to labeling, partially expanded (∼2×) hydrogels were blocked in 3% bovine serum albumin (BSA) and 0.01% Triton X-100 in 1× PBS. All samples were labeled via indirect immunofluorescence or covalently reactive small-molecule fluorescent dyes (e.g., NHS-ester conjugates). Specifically, samples were concurrently immunolabeled with mix of previously centrifuged primary antibodies raised against distinct host species (**Table S1**) diluted at 1:50 – 1:1000 (v/v) in staining buffer, which consists of PBST [0.01% Triton-X in PBS] supplemented with 1% BSA, for 2.5 h at 37 °C. After primary antibodies incubation, samples were washed three times with an excess of PBST for 20 min each. Hydrogels were then protected from light and incubated in staining buffer containing a mix of highly cross-adsorbed secondary antibodies (**Table S2**), each diluted 1:500 (v/v), for 2 h at 37 °C, followed by a repetition of the washing steps. For nuclear staining, DAPI (0.3 µM, Thermo Fisher Scientific Cat# 62248) was used. For antibody stripping, hydrogels were incubated in 3 mL of denaturation buffer supplemented with 100 mM BME at 95 °C for 45 min and washed for at least 1 h with an excess of 1× PBS to remove detached antibodies. Blocking and immunostaining were followed by antibody stripping and repeated over eleven iterative rounds (**Table S6**). In each round, the microtubule network was relabeled to provide a consistent structural reference for multi-target image registration.

### Pre-imaging gel processing

To enable iterative, volumetric imaging of defined regions of interest, expanded gels were oriented with the cellular surface facing downward and trimmed into irregular shapes matching the dimensions of the imaging dish (u-Slide 8 Well High Glass Bottom, Idibi Cat #80807). This approach minimized lateral drift and ensured consistent repositioning across imaging rounds. After removing excess water, hydrogels were placed directly onto the glass surface without Poly-D-lysine or other adhesives, as these fragile gels can tear or deform when mechanically mounted. For extended imaging sessions, gels were optionally secured by affixing the edges to a glass-bottom dish (MatTek Cat #P35G-1.5-14-C) using a small amount of hot glue applied outside the field of view

### Spinning Disk Confocal Microscope

Cells shown in **Figure 1c, Supplementary Fig. 1, 4, and 6-9** were imaged using a Nikon CSU-W1 SoRa inverted spinning disk confocal microscope (Nikon Instruments), equipped with a SoRa super-resolution module. The system features a dual-disk scan head, a piezo Z-drive for rapid volumetric acquisition, a motorized XY stage for multi-position imaging and tiling, and Nikon’s Perfect Focus System to maintain focus stability throughout extended acquisitions. Excitation was provided by 405, 445, 488, 514, 561, and 640 nm laser lines. Emitted fluorescence was detected using a Hamamatsu ORCA-Fusion sCMOS camera (C14440-20UP). Imaging was performed using multiple infinity-corrected objectives from Nikon, covering a broad range of magnifications. Depending on the application, an air-immersion Plan Fluor λ 20×/0.75 objective (working distance (WD): 1.0 mm, MRD00205) was used for larger field-of-view imaging. Higher-resolution imaging was performed using oil-immersion objectives, including a Plan Fluor 40×/1.30 objective (WD: 0.24 mm, MRH01401), a Plan Apo λ 60×/1.40 objective (WD: 0.13 mm, MRD01605), and a Plan Apo λ 100×/1.45 objective (WD: 0.13 mm, MRD01905). All objectives were designed for a standard cover glass thickness of 0.17 mm, providing a high numerical aperture for subcellular imaging. Z-stacks were acquired using step sizes of 0.9 µm, 0.3 µm, or 0.2 µm, as appropriate for the axial resolution and sampling requirements.

### Oblique Plane Microscope

A custom-built oblique plane microscope (OPM) was used for imaging cells in **Figures 1d–k, i-k, 2, Supplementary Figures 2, 3, 5, and 10.** The OPM system was constructed in an inverted geometry and optimized for high-resolution imaging of aqueous specimens^15^. The sample was illuminated obliquely at 45° using a water immersion objective (25×, NA 1.1, MRD77220, Nikon). Fluorescence was collected through the same primary objective in an epi-fluorescence format and relayed to a secondary objective (20×, NA 0.8, UPLXAPO20X, Olympus). The emission was then redirected at a 45° angle by a custom glass-tipped tertiary objective (AMS-AGY v2, 54-18-9, Applied Scientific Instrumentation) and projected onto an sCMOS camera (ORCA-Flash4.0, C13440-20CU, Hamamatsu) via a tube lens (AC508-300-A-ML, Thorlabs). The effective isotropic pixel size of 150 nm after shear/rotation was previously validated empirically by imaging calibration specimens of known dimensions (e.g., fluorescent bead samples and stage micrometer/grid standards) and confirming the expected scaling in all axes^15^. Excitation was provided by 405, 488, 561, and 640 nm laser lines. The system was controlled with custom LabView software interfaced through National Instruments I/O hardware (National Instruments). Z-stacks were acquired using step sizes of 0.21 µm or 0.32 µm.

### Imaging Parameters

Unless otherwise noted, all cells were volumetrically imaged in their 4.2× expanded state to capture the full extent of their three-dimensional structure. All detailed acquisition settings, including objective types, laser powers, emission filters, Z-step sizes, and other camera configurations, are listed in **Table S8**.

### Computing, Image Analysis, and Image Registration

All computational analyses were performed on a SLURM-managed high-performance computing cluster. Hardware specifics are not critical to the results; for completeness, analyses were run on Red Hat Enterprise Linux 9.4 nodes with 512 GB RAM and dual Intel Xeon Gold 6354 CPUs (72 logical CPUs). Image-processing, registration, and analysis routines were implemented in Python (version ≥3.11). JupyterLab was used for interactive development, while long-running registration and analysis pipelines were executed as SLURM scripts. Images from each staining cycle were registered using the Advanced Normalization Tools Python interface (ANTsX/ANTsPy)^19^. Initial alignment was performed with the ‘TRSAA’ transformation method, which sequentially applies Translation, Rigid, Similarity, and two Affine transformations to achieve a high-quality affine mapping. This step was configured with multi-resolution iterations set to (1000, 500, 250, 100) to robustly converge. To further refine the alignment and correct residual non-linear deformations, we performed an elastic (non-linear) registration step using a smoothly varying deformation field. This procedure applied moderate flow smoothing and minimal total-field regularization and was executed at a single resolution level to ensure that only local, fine-scale adjustments were introduced. Additional dependencies included volumetric file I/O using tifffile, image processing via scikit-image and SimpleITK, and plotting using matplotlib and seaborn. All custom image-processing and registration routines are publicly available on GitHub (https://github.com/TheDeanLab/clearex), with a versioned and archived release to be deposited on Zenodo upon manuscript acceptance.

### Highly Multiplexed Image Processing

The raw image volumes were geometrically transformed to a standard orientation via shearing and rotation operations implemented in MATLAB as previously reported^20^. Registered z-stacks were imported into ImageJ (v1.54r) as separate grayscale images and merged into a single hyperstack, with each slice assigned as an individual channel. All channels were pseudo-colored by applying individual lookup tables (LUTs), and linear intensity ranges were manually adjusted using Brightness/Contrast. The final images were exported as 16-bit TIFF files for **Figure 2a** preparation. The 20-plexed visualization of cellular structures (**Fig. 2b**) was performed in napari (0.6.6), where channel-specific colormaps were individually applied. Opacity, intensity scaling, and gamma correction were manually adjusted to enable simultaneous visualization of dense cellular structures, and images were exported at full spatial resolution for downstream figure assembly.

### Analysis of Staining Intensity

To evaluate fluctuations in staining intensity across imaging rounds, peroxisomes and nuclei were repetitively labeled, imaged, and stripped. Imaging was performed using a spinning disk confocal microscope with constant acquisition settings between rounds. Three-dimensional image stacks were converted to maximum intensity projections, and each round was registered to the initial round (chosen as reference) using the TRSAA algorithm implemented in ANTsX/ANTsPy. Regions of interest of identical size were then selected from each cell within the field of view, and the background-corrected mean intensity was extracted for quantitative comparison across cycles.

### Nuclei Size Quantification

For nuclear area measurements (**Figure S4e and g**), nuclei were stained with SYTOX Green (0.167 µM, Invitrogen Cat# S7020) during each imaging round. Nuclei were imaged before and after full (4.2×) expansion using a spinning disk confocal microscope, ensuring that each Z-stack encompassed the entire nuclear volume. Maximum intensity projections were generated in ImageJ (v1.54r) to produce 2D representations suitable for area quantification. Segmentation of individual nuclei was performed using Cellpose with default nuclei-segmentation parameters. Nuclei area was calculated by summing the pixels within each nucleus mask and converting pixel counts to physical units using the native pixel dimensions of the raw images. Measurements are reported in µm². For estimation of nuclear “diameter,” we report an operational linear size estimate measured along a single user-defined axis across each nucleus (i.e., a one-dimensional line measurement rather than a fitted equivalent diameter). Nuclear areas (**Fig. S3e**) and diameters (**Fig. S3g**) were visualized in RStudio (R version 4.4.0) using the dplyr, tidyr, ggplot2, and ggpubr packages.

### Mitochondria Detection

Images were preprocessed with background subtraction and intensity rescaling. Candidate puncta were detected using Difference-of-Gaussians algorithm implemented in Skikit-image. The nominal object size was parameterized by a Gaussian-equivalent FWHM of 5 pixels, with a multi-scale search. Detection thresholds were set on a per-image basis, reflecting round-dependent intensity differences. Detected object coordinates were stored in row/column coordinates for subsequent pairing.

### Nearest-Neighbor-Based Target Registration Error Estimate

To estimate registration accuracy independently of the reference channel, we quantified a target-registration-error (TRE) proxy by (i) registering imaging rounds using the microtubule channel and (ii) measuring residual point-to-point misalignment on mitochondrial puncta, which were not used to compute the registration transforms. To expedite analysis, volumetric tiff stacks were reduced to 2D maximum-intensity projections. Five staining rounds were analyzed, and registration was performed pairwise using round 1 as the fixed reference and rounds 2–5 as moving images. This included the ANTsPy TRSAA transform, which applies a pyramidal sequence of transforms to obtain a robust mapping between rounds. The resulting transform was then applied to both microtubules and mitochondria. To correct residual local distortions, a deformable registration was performed on the TRSAA-aligned microtubules using a nonlinear registration (type “SyNOnly”). For mitochondria, we applied the affine TRSAA transform followed by the SyN deformation field; in ANTsPy this was implemented using the standard transform list ordering (SyN transforms listed before TRSAA) when resampling the original moving mitochondria image.

Mitochondrial puncta were detected independently in the fixed mitochondria MIP and in each transformed mitochondria MIP under both transform conditions. To estimate a TRE proxy, puncta were paired between the fixed and transformed mitochondria images using mutual nearest-neighbor (MNN) matching. Specifically, a k-d tree was constructed for each point set, each moving point was assigned its nearest fixed neighbor, and each fixed point its nearest moving neighbor. Pairs were retained only if the association was mutual (moving to fixed and fixed to moving). Pairing was evaluated with multiple distance gates (unlimited, 10 px, 5 px, 3 px) to characterize sensitivity to the matching threshold. For each condition and round, we summarized the distribution of paired distances by mean, median, standard deviation, 95th percentile (P95), maximum, and RMS distance. Distances were corrected for specimen expansion using an effective pixel size of 38.7 nm.

### Descriptor– and RANSAC-based Target Registration Error Estimate

To obtain a more conservative estimate of residual misalignment that is not driven by local, ambiguous nearest-neighbor assignments, we complemented mutual nearest-neighbor pairing with a descriptor– and RANSAC-based landmark matching approach. Here, “globally consistent” refers to correspondences that are jointly explainable by a single best-fit similarity transform (translation/rotation/scale) between rounds, rather than independent local pairings. Mitochondrial puncta coordinates (detected as described above) were treated as candidate landmarks for feature matching between imaging rounds. For each pai wise comparison, maximum-intensity projections were used, and images were cropped to a common overlapping field of view when necessary to ensure identical dimensions prior to matching. Binary BRIEF descriptors were computed at mitochondrial landmark locations, and putative correspondences were identified by matching descriptors using Hamming distance. Because descriptor matching can include spurious pairs, RANSAC was used to robustly fit a similarity-transform model and reject outliers, retaining only inlier landmark pairs consistent with a single global transform. The similarity transform was then refit using all inliers. To avoid reporting an optimistically biased fit error, we computed a cross-validated TRE-like proxy by repeatedly fitting the similarity transform on a subset of inlier correspondences and evaluating residual distances on held-out inliers; the root mean square of these held-out residuals was used as the final descriptor-based registration accuracy estimate.

### Antibody Signal Distribution Analysis

Cells and nuclei were individually segmented with CDC42 as a diffuse cytoplasmic marker, and DAPI as a nuclei marker using u-Segment3D^21^ and pretrained Cellpose^22^ 2D segmentation models. The distance transform was computed using u-Unwrap3D^23^ and is the Fast-Marching Method solution to the Eikonal equation with the nuclei mask as a source term, and the cell mask as a boundary condition^21^ We normalize the distance transform by dividing by the maximum value after masking out the cell exterior and nuclear interior regions. The normalized distance, 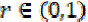 was then equipartitioned into 50 intervals. The signal distribution of each antibody was constructed as the median voxel intensity in each interval. To group signal distributions, we constructed the pairwise Pearson correlation matrix and applied hierarchical clustering to the matrix with a Euclidean metric and Ward linkage.

## SUPPLEMENTARY FIGURE CAPTIONS

**Figure S1.**
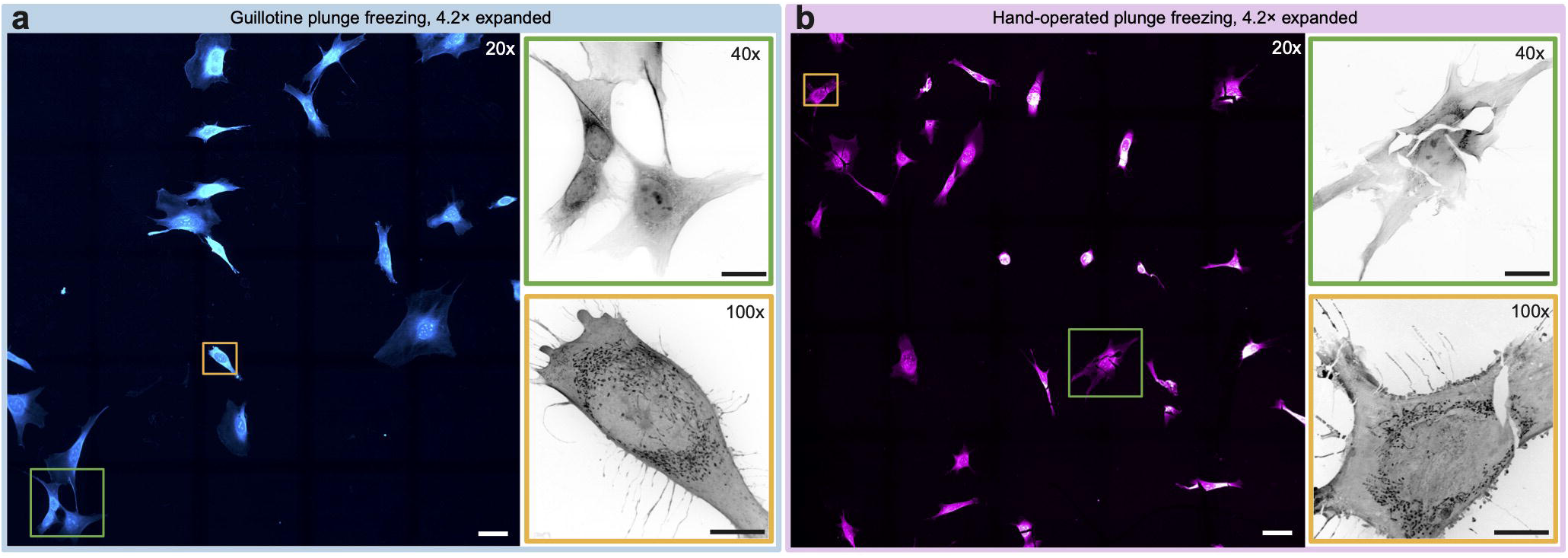
Impact of Freezing Kinetics on Cellular Integrity. Representative spinning disk confocal images of 4.2× expanded and N-hydroxysuccinimide ester stained MEF cells subjected to: (a) rapid guillotine plunge freezing and (b) hand operated freezing. Insets squares highlight selected cells shown at higher magnification. Cracks and ruptures in cellular architecture were more frequently observed following hand-operated freezing, as seen in panel b. Imaging was performed using 20×, 40×, and 100× objectives. Maximum intensity projections are shown. Scale bars: 20 µm (corrected for the expansion factor).

**Figure S2.**
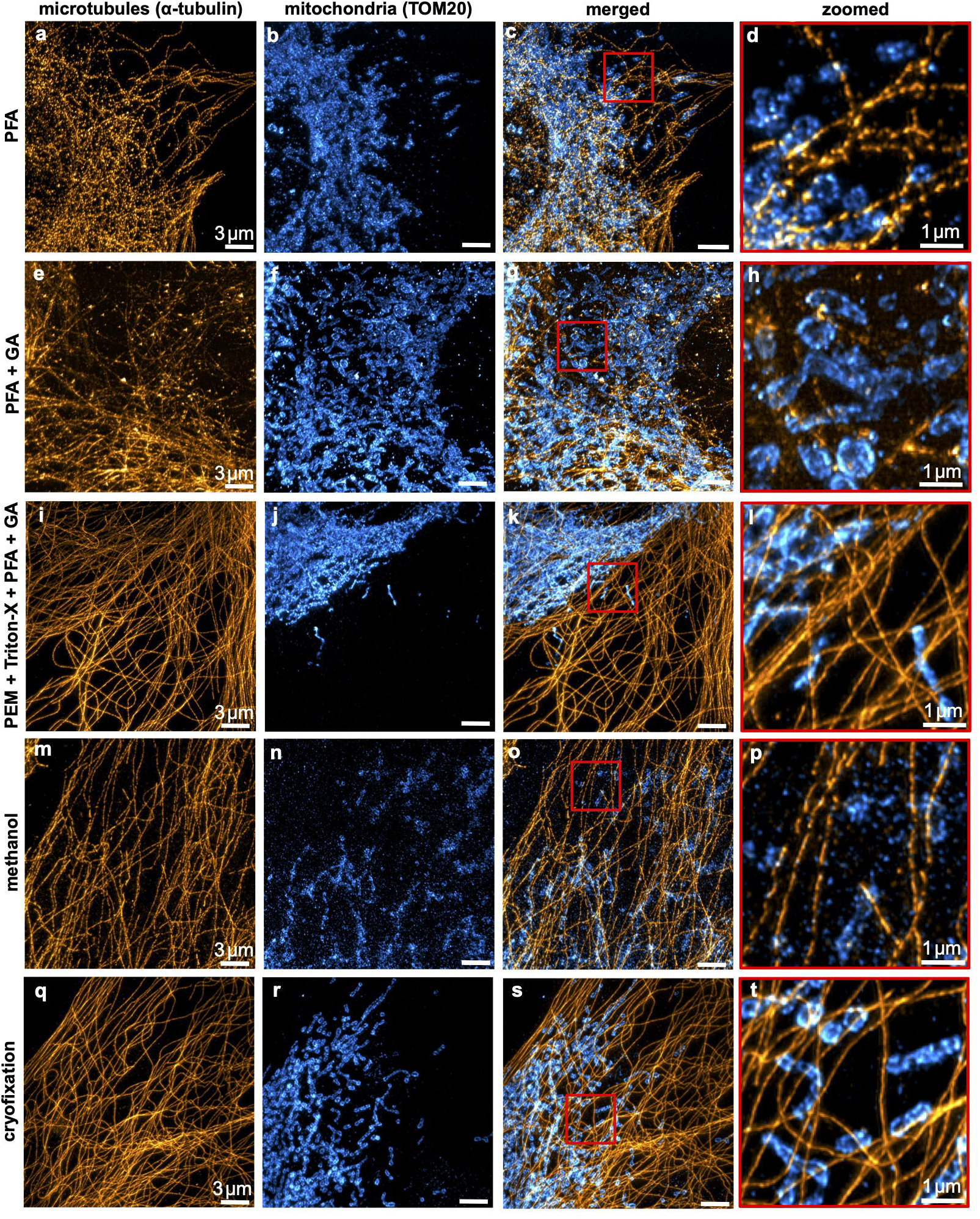
Comparison of Fixation Methods. Deconvolved, maximum intensity projections of microtubules (anti-α-Tubulin, Sigma-Aldrich Cat# T9026, RRID:AB_477593) and mitochondria (anti-TOM20, Proteintech Cat# 11802-1-AP, RRID:AB_2207530) in 4.2× expanded MEF cells subjected to: (a–d) 4% paraformaldehyde (PFA) fixation, (e–h) 4% PFA + 0.2% glutaraldehyde (GA) fixation, (i–l) cytoskeleton-preservation-optimized fixation^18^, (m–p) 100% methanol fixation, (q–t) and cryofixation by guillotine-style plunge-freezing. Red squares in merged images correspond to zoomed regions of interest shown in (c, g, k, o, s). Images acquired with a custom-built oblique plane microscope. All scale bars are corrected for the expansion factor. While traditional fixation methods are effective for select structures, cryofixation offers the most comprehensive preservation of native cellular architecture, effectively maintaining the native architecture of cytoskeletal compartments as well as membrane-bound organelles. Methanol, for example, is commonly used for cytoskeletal preservation but can induce depolymerization of microtubules and actin filaments. Aldehyde-based fixatives such as PFA form covalent crosslinks that may mask epitopes and necessitate additional antigen retrieval steps. In contrast, cryofixation preserves proteins in their native state without chemical modification, resulting in superior epitope accessibility and antigenicity for high-resolution immunofluorescence imaging.

**Figure S3.**
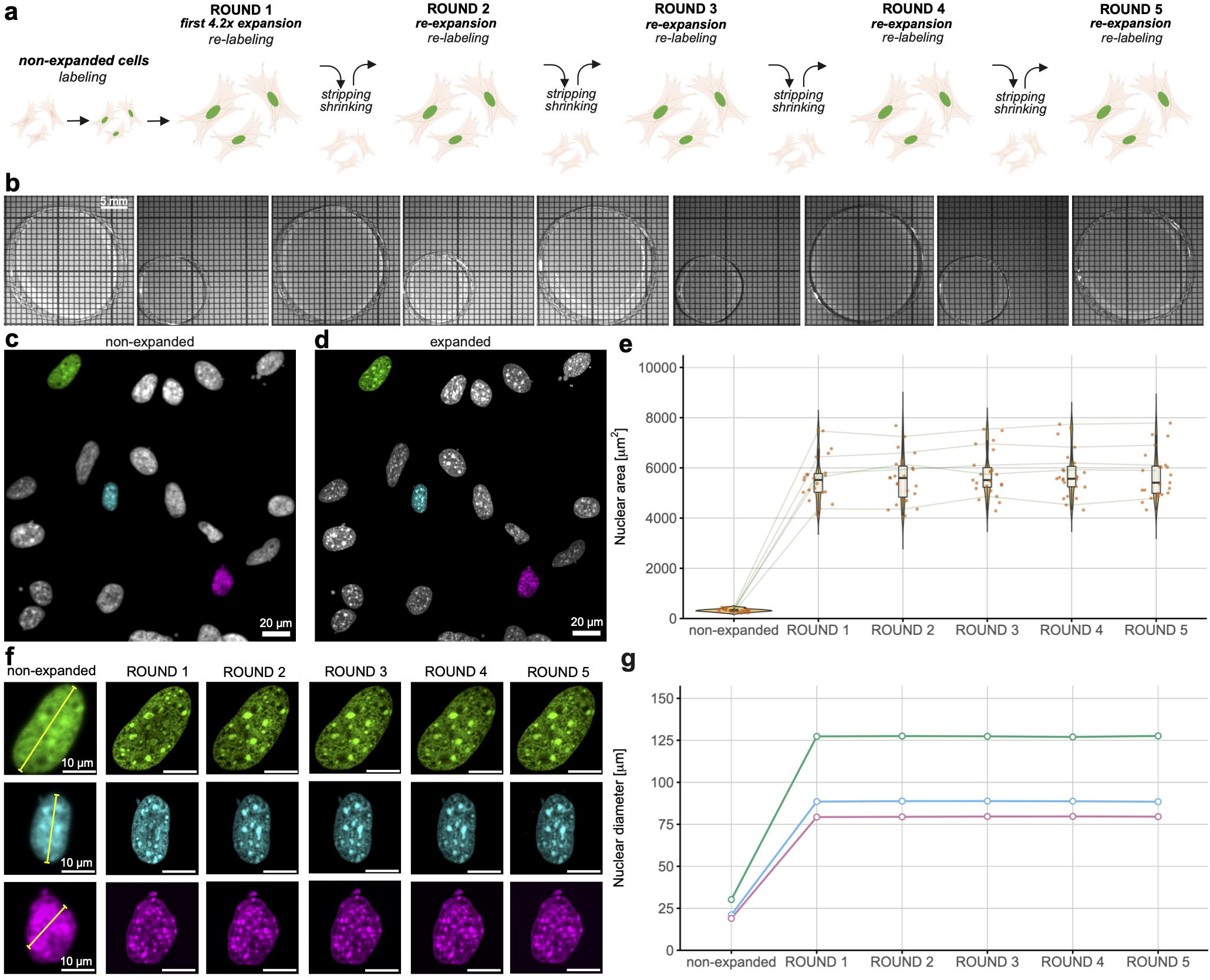
Enhanced resolution in expanded cells visualized by Cy-ExM. (a) Non-expanded and (b) expanded MEF cells stained for DNA (SYTOX Orange, Invitrogen Cat# S11368) and the outer mitochondrial membrane (anti-TOMM20, Abcam Cat# ab232589, RRID:AB_3065091). Representative but distinct cells are shown in each condition. Cells were imaged volumetrically using an oblique plane microscope. Deconvolved maximum intensity projections are shown for standard immunofluorescence (a) and for cells processed with a single round of the Cy-ExM workflow (b). Insets highlight representative subcellular features, demonstrating improved spatial resolution and structural detail following 4.2× expansion. Scale bars: 10 µm (overview), 1 µm (insets), corrected for the expansion factor.

**Figure S4.**
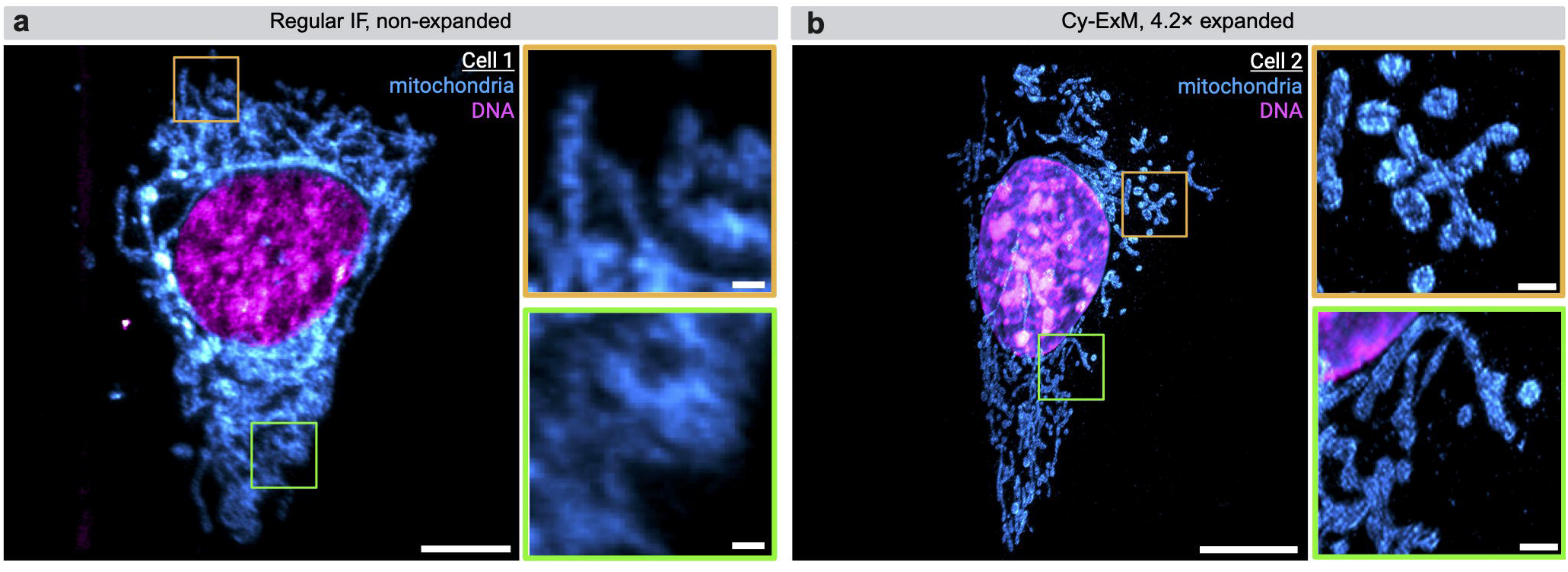
Validation of expansion consistency and iterative labeling workflow. (a) Schematic illustrating the iterative labeling workflow used in Cy-ExM. (b) Macroscopic gel conditioning across rounds. Representative images of the same hydrogel alternating between full expansion in deionized water and partial shrinkage in 1× PBS during staining. The apparent ∼2× change in gel diameter reflects this reversible water/PBS conditioning and is shown to illustrate handling during iterative cycles; it is not the 4.2× cellular expansion factor used for imaging. (c–d) Spinning disk confocal images of the same population of MEF cells stained for DNA (SYTOX Green, Invitrogen Cat# S7020), acquired before (c) and after (d) full 4.2× expansion. (e) Quantification of expansion repeatability across labeling rounds by measuring nuclear area in the fully expanded state. Green lines link corresponding nuclei across rounds to visualize consistency. (f) Example of three randomly selected nuclei tracked and iteratively imaged over five expansion rounds; yellow lines denote positions used for diameter measurements. (g) Measured nuclear diameters of the three nuclei across rounds, confirming consistent expansion. Line colors correspond to the artificially assigned nuclear colors shown in (f). The 4.2× linear expansion factor was calculated from nuclear area as the square root of the area in the expanded state divided by the area in the unexpanded state. All imaging was performed under identical acquisition conditions for each round. All scale bars are corrected for the expansion factor.

**Figure S5.**
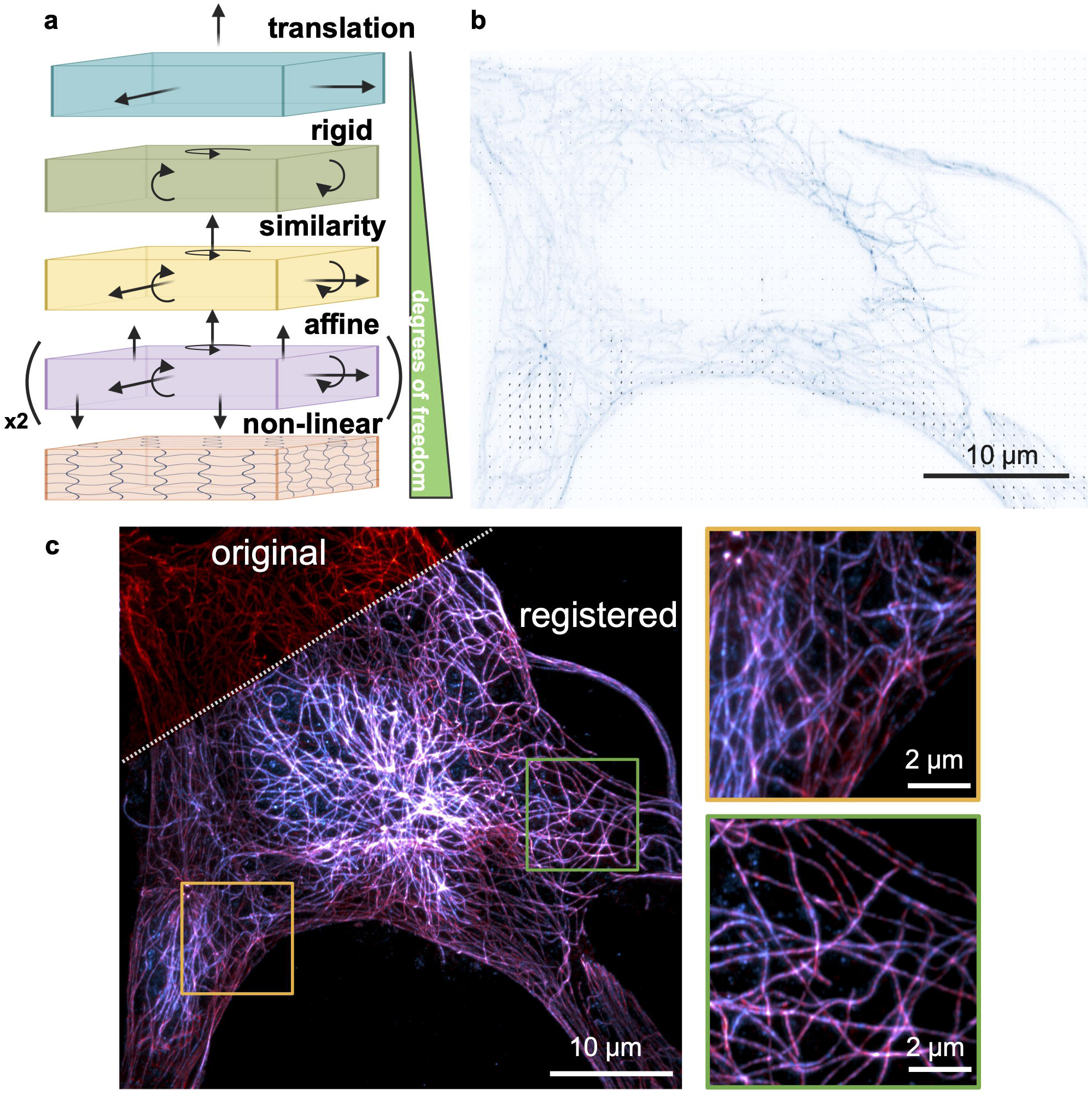
Registration Strategy, Deformation Quantification, and Overlay of Registration Results. **(a)** Schematic of the registration pipeline. Samples were sequentially aligned using translation, rigid body, similarity, and affine transforms, followed by a non-linear warp registration to correct for local distortions. **(b)** Vector plot illustrating local deformations from the warp field for a representative 2D section of the image. **(c)** Result of the registration routine enabling precise alignment of the microtubule network across independent labeling rounds. Original image in top left corner (first staining, red) and registered overlay (first and second staining, red + cyan) are shown, with inset squares. All scale bars were corrected for the expansion factor.

**Figure S6.**
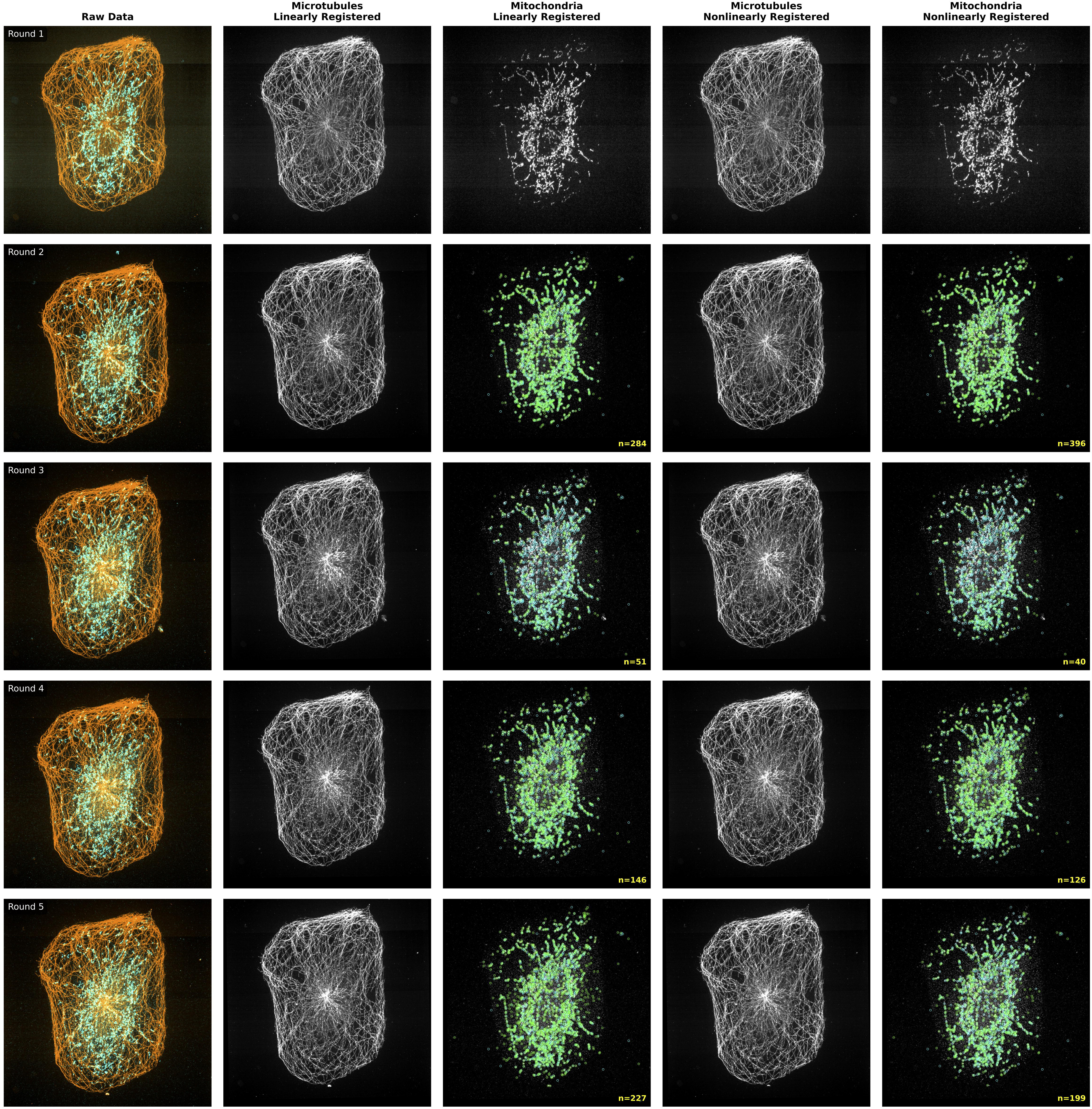
Multi-round registration accuracy assessed by microtubule-guided alignment and independent mitochondrial landmark matching. Five imaging rounds are shown (rows), with each column depicting a successive stage of the registration workflow. Left: raw overlays of microtubules (orange) and mitochondria (cyan). Second column: microtubule channel after linear registration to round 1. Third column: mitochondria channel after applying the linear transform, with detected mitochondrial puncta overlaid; cyan and lime circles denote puncta detected in the fixed (round 1) and transformed moving images, respectively. Fourth column: microtubule channel after both linear and nonlinear registration. Fifth column: mitochondria channel after applying the combined linear and non-linear transforms, with matched puncta displayed as in the third column. Round 1 serves as the fixed reference and is therefore identical across conditions. Imaging was performed with a spinning disk confocal microscope at 40x magnification.

**Figure S7.**
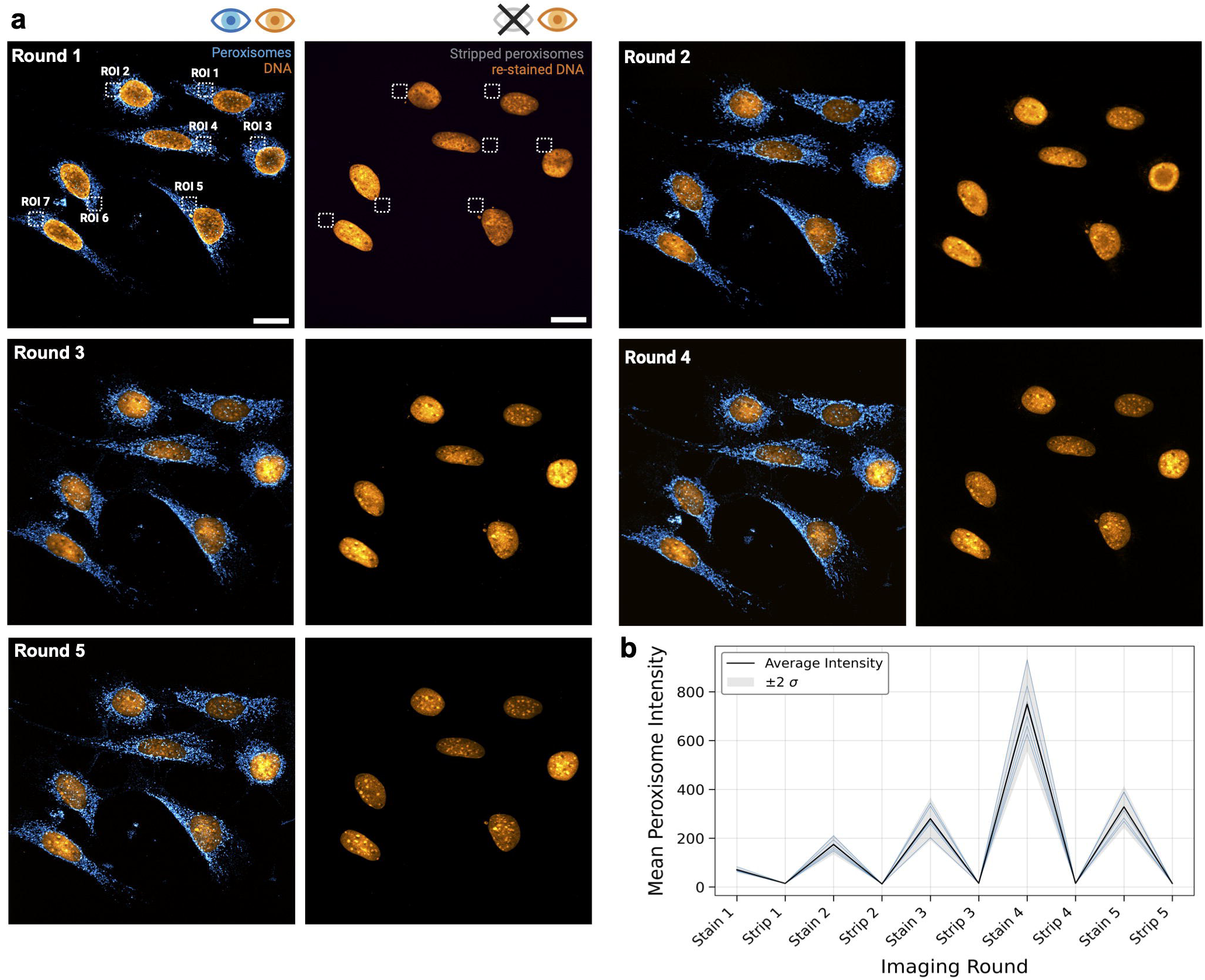
Iterative peroxisome imaging validates registration accuracy and stripping efficiency. **(a)** 4.2× expanded MEF cells stained for DNA (SYTOX Green, Invitrogen Cat# S7020) and peroxisomes (anti-PMP70, Thermo Fisher Scientific Cat# PA1-650, RRID:AB_2219912) were iteratively imaged across five rounds of the Cy-ExM workflow. Shown is the same colony of cells in the labeled state (left) and after successful antibody removal (right). For each round, identical labeling conditions were used (antibody concentration, incubation time, and temperature), followed by overnight expansion in deionized water. In the image from Round 1, dashed-line squares indicate the regions of interest (ROIs) where fluorescence signal intensity was consistently measured in each round. Imaging was performed on a spinning disk confocal microscope using a 20× objective, with unchanged acquisition settings. For peroxisomes: 640 nm laser at 60 mW and 200 ms exposure was applied every time, ensuring reproducibility for average signal intensity quantification shown in (b). Scale bars: 20 μm (corrected for the expansion factor).

**Figure S8.**
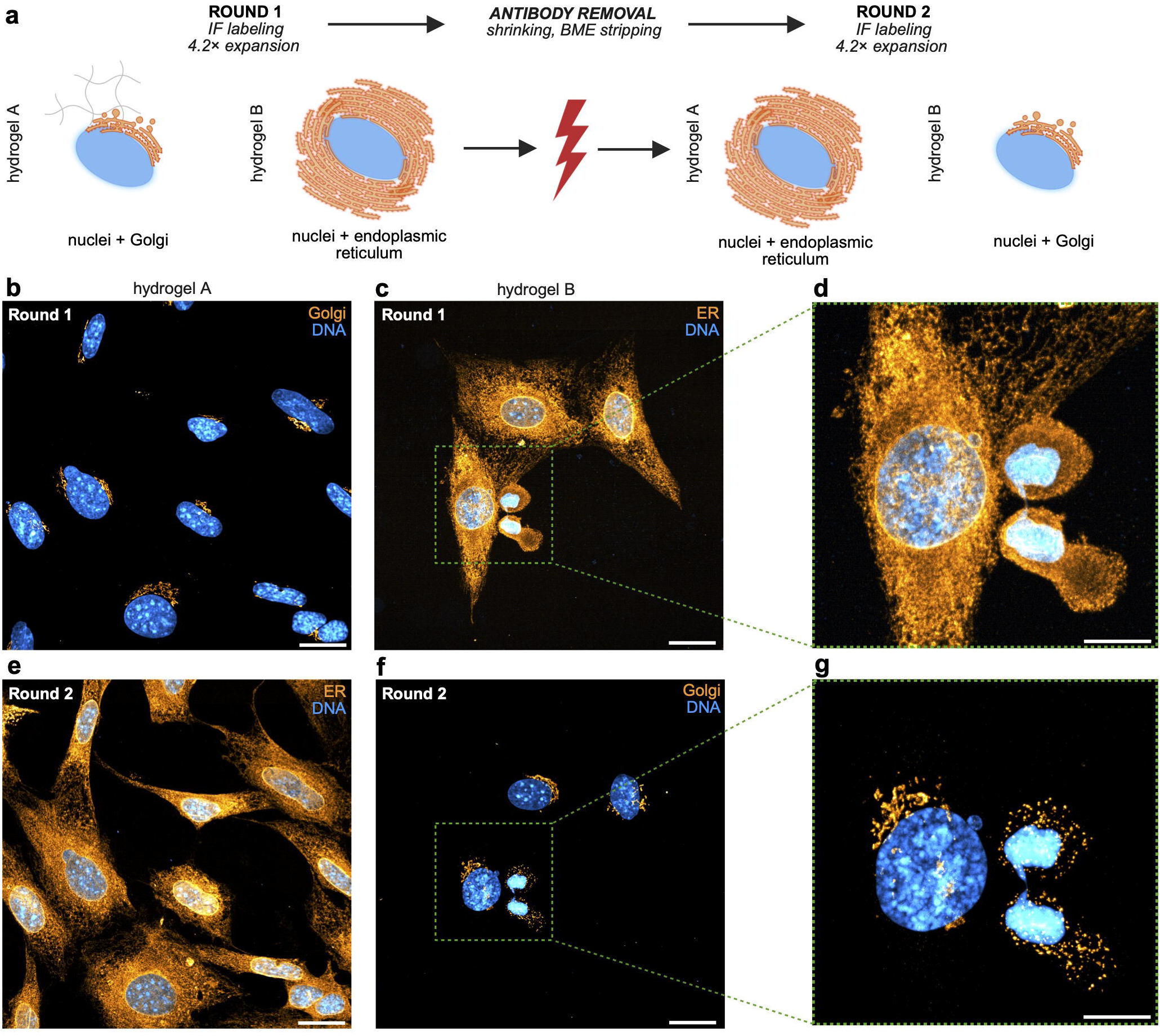
Cycle permutation confirms antigen stability across rounds. (a) Schematic of the permutation experiment workflow. (b) Cells embedded in hydrogel A were expanded and stained for DNA (DAPI, Thermo Scientific Cat# 62248) and the Golgi apparatus (anti-GOLGA2/GM130, Proteintech Cat# 11308-1-AP, RRID:AB_2115327), while (c) cells in hydrogel B were simultaneously stained for DNA (DAPI) and the endoplasmic reticulum (ER; anti-calnexin, Abcam Cat# ab22595, RRID:AB_2069006). Following β-mercaptoethanol (BME) treatment to remove antibodies, the hydrogels were re-stained in the opposite configuration: hydrogel A for DNA and ER, and hydrogel B for DNA and Golgi, using the same antibodies and labeling conditions. Insets in (d) and (g) highlight a MEF undergoing late cytokinesis (telophase-to-G1 transition), a stage characterized by organelle reorganization and nuclear reassembly during daughter cell separation. The cell contains two nascent daughter nuclei connected by a persistent cytoplasmic bridge, consistent with incomplete abscission. (d) reveals extensive ER continuity across the bridge, suggesting active ER remodeling, while (g) shows a fragmented Golgi apparatus distributed around the daughter nuclei, consistent with Golgi vesiculation during mitotic exit. All scale bars are corrected for the expansion factor: 20 µm (b, c, e, f) and 10 µm (d, g). Images were acquired using a spinning disk confocal microscope with a 20× objective under identical acquisition settings.

**Figure S9.**
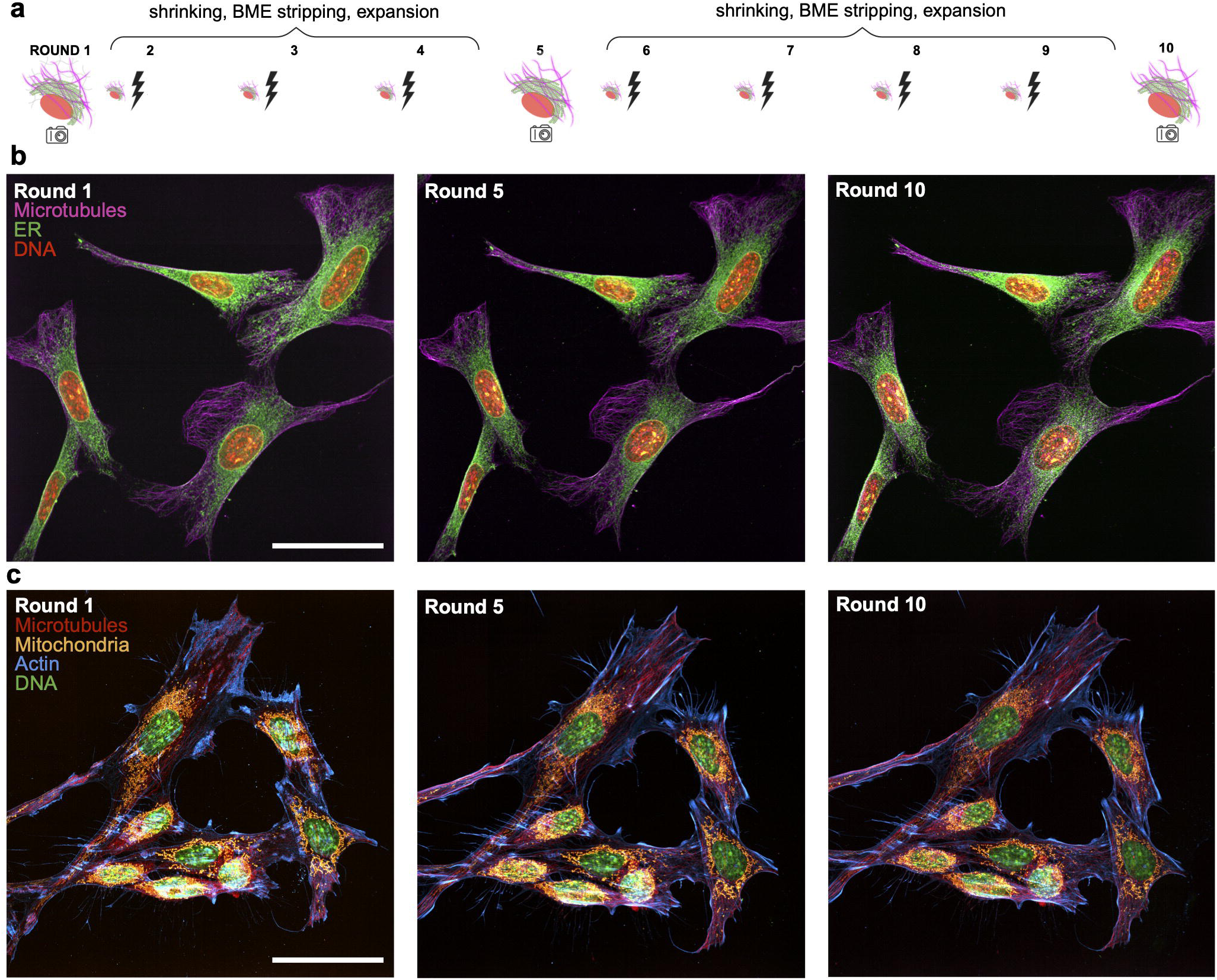
Benchmarking Cy-ExM across repeated labeling cycles. (a) Schematic of the experiment: hydrogels with MEF cells were subjected to 10 Cy-ExM cycles, each including: 1 h blocking, 2.5 h incubation in primary antibody staining solution, 1 h of washing, 2 h incubation in secondary antibody staining solution, followed by 2 h of physical expansion in water, and 45 min of β-mercaptoethanol treatment in 95 °C. Two hydrogels were independently stained: first (b) for microtubules (anti-α-Tubulin, Sigma-Aldrich Cat# T9026, RRID:AB_477593), endoplasmic reticulum (anti-calnexin, Abcam Cat# ab22595, RRID:AB_2069006) and DNA (DAPI, Thermo Scientific Cat# 62248), while the second gel (c) was stained for microtubules (anti-α-Tubulin, Selleckchem Cat# F1566), mitochondria (anti-HSP60, Abcam Cat# ab46798, RRID:AB_881444), actin filaments (anti-β-actin, Sigma-Aldrich Cat# A1978, RRID:AB_476692) and DNA (DAPI). Hydrogels were imaged every 5 rounds using a spinning disk confocal microscope with a 20× objective under identical acquisition settings. Scale bars: 50 µm (corrected for the expansion factor).

**Figure S10.**
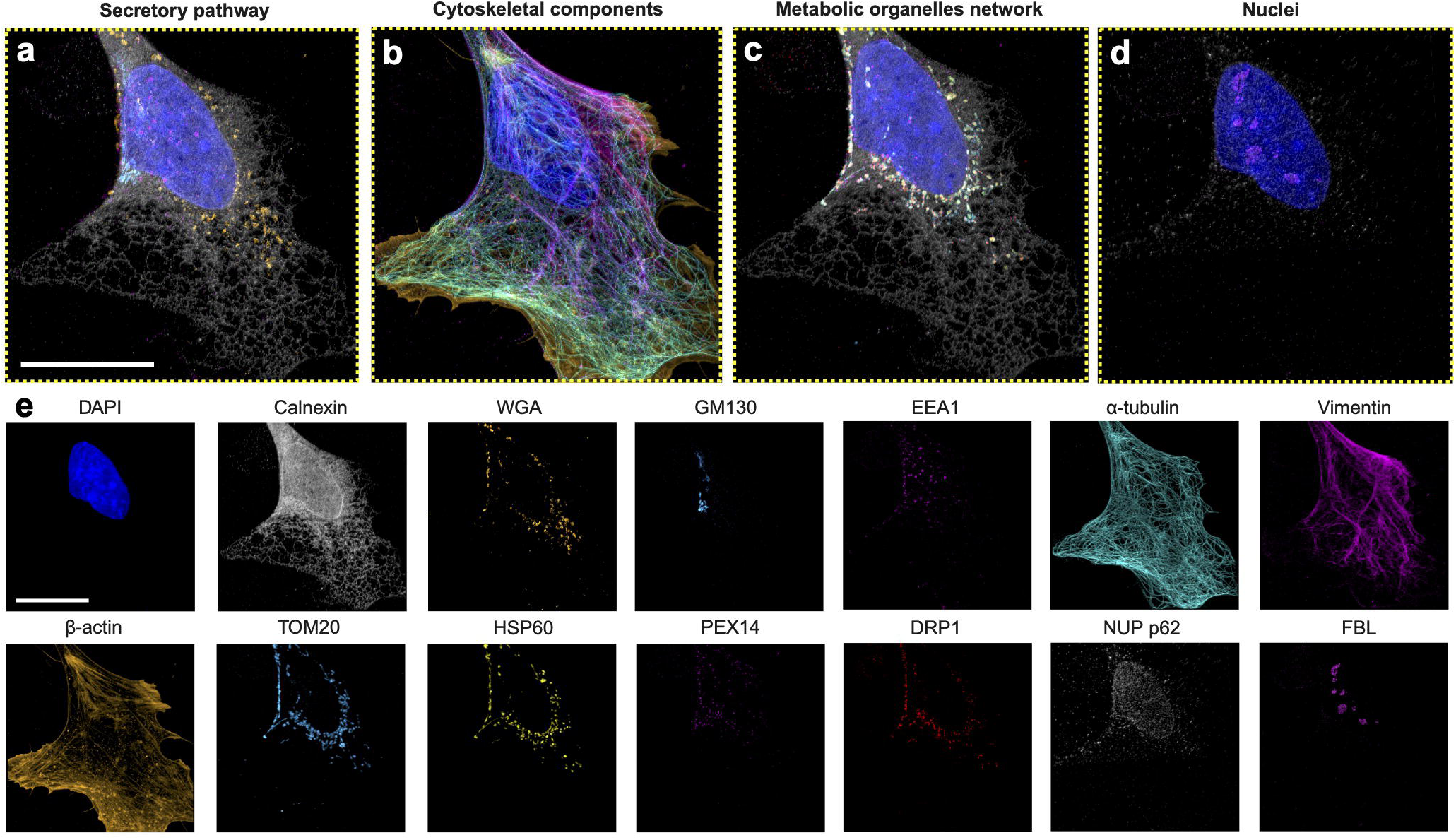
Fourteen-plex Cy-ExM imaging of subcellular organization in a MEF cell. (a–d) Maximum-intensity projections (MIPs) from the same cell with targets grouped by biological function: (a) secretory pathway, (b) cytoskeletal components, (c) metabolic organelle network, and (d) nuclear organization. (e) Corresponding single-channel MIPs for each of the 14 targets shown individually; colors in (e) match the merged images corresponding to the functional classes in (a–d). All samples were processed using the Cy-ExM workflow and imaged volumetrically using an oblique plane microscope. Scale bar: 5 µm (corrected for the expansion factor).

## SUPPLEMENTARY MOVIE CAPTIONS

**Supplementary Movie 1. Three-dimensional volume rendering of peroxisomes, microtubules, and nuclei.** Data were deconvolved and geometrically reoriented to correct the oblique plane microscopy imaging geometry (shear/deskew) and rotated into the standard viewing perspective of an inverted microscope (see **Methods**). Volumetric imaging was performed using an oblique plane microscope. Scale bar: 5 µm.

**Supplementary Movie 2. Z-stack rendering of the registered microtubule network across two independent immunolabeling rounds.** The overlay of the first staining (red) with the registered second staining, after antibody stripping (cyan) demonstrates the accuracy of the registration routine and capability to repeatedly target the same cellular component. Imaged volumetrically using an oblique plane microscope. Scale bar: 5 microns.

**Supplementary Movie 3. Three-dimensional visualization of a 20-plex Cy-ExM dataset.** Three-dimensional rendering of 20 distinct cellular targets from a single cell, grouped according to their primary biological function, including nuclear/nucleolar components, mitochondria, secretory pathway organelles, endomembrane trafficking compartments, cytoskeletal structures, and adhesion-related proteins. The volume is displayed as an attenuated maximum-intensity projection with additive blending to highlight spatial relationships among molecular assemblies throughout the cellular volume. Colored bars adjacent to each label indicate the per-channel intensity look-up table used for visualization. Data were imaged volumetrically using an oblique plane microscope.

**Supplementary Movie 4. Two-dimensional Z-flythrough of a 20-plex Cy-ExM dataset grouped by biological function.** A 20-target Cy-ExM dataset from a single cell is shown as a sequence of 2D optical sections. Targets are grouped by primary biological function (nuclear, mitochondria, secretory pathway organelles, endomembrane trafficking, and cytoskeletal structures) to facilitate interpretation of subcellular organization across depth. The current Z position is annotated in the upper-left corner of the movie. Colored bars adjacent to each label indicate the per-channel intensity look-up table used for visualization. Data were acquired volumetrically using an oblique plane microscope.

## SUPPLEMENTARY TABLES

**Table S1.**
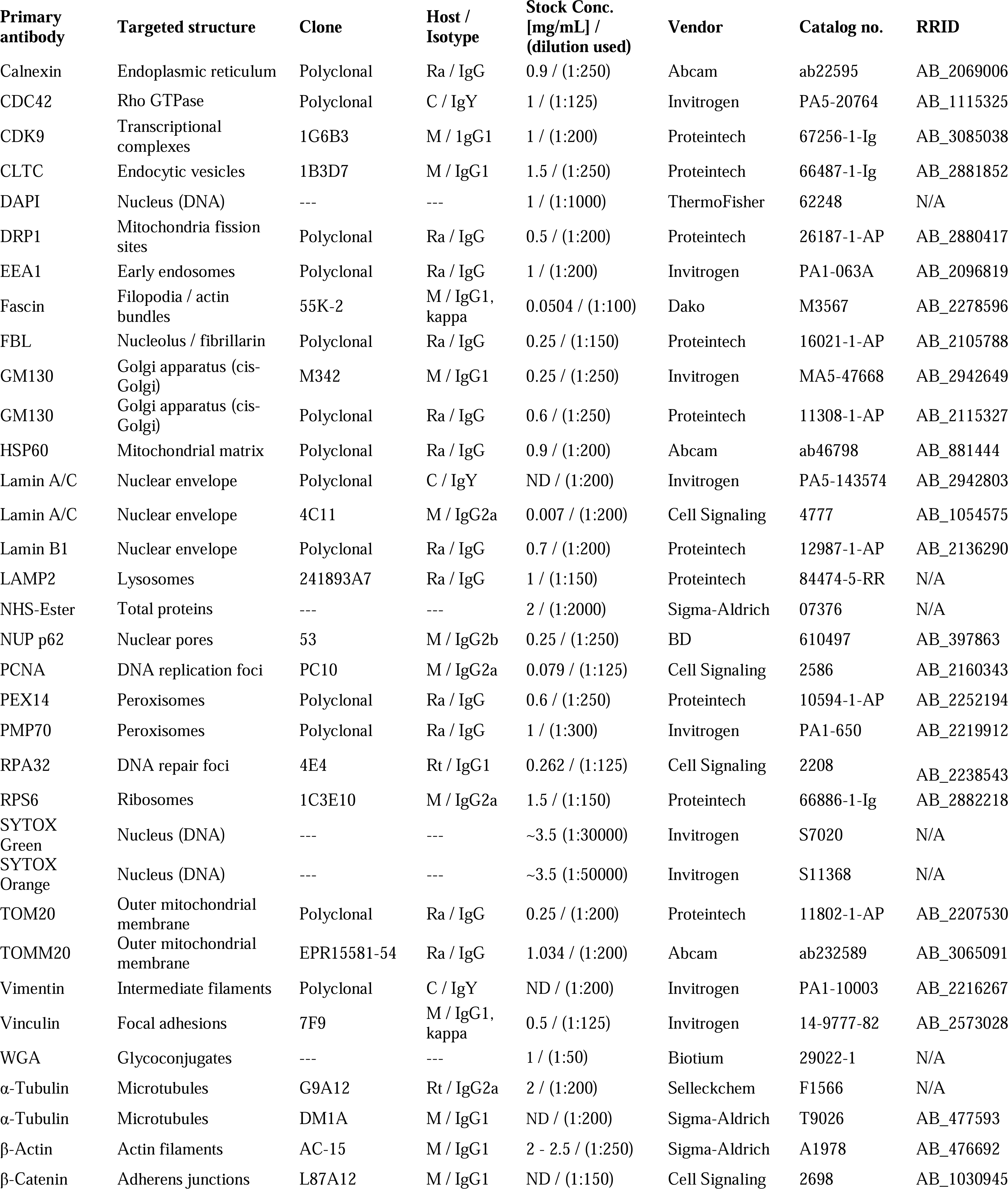
Primary antibodies and dyes used in this study. The table lists targets, clone information, host species and isotype, stock concentration, working dilution, vendor, and catalog number for each reagent. Abbreviations: host—Ra, rabbit; C, chicken; M, mouse; Rt, rat; stock concentration—ND, not determined.

**Table S2.**
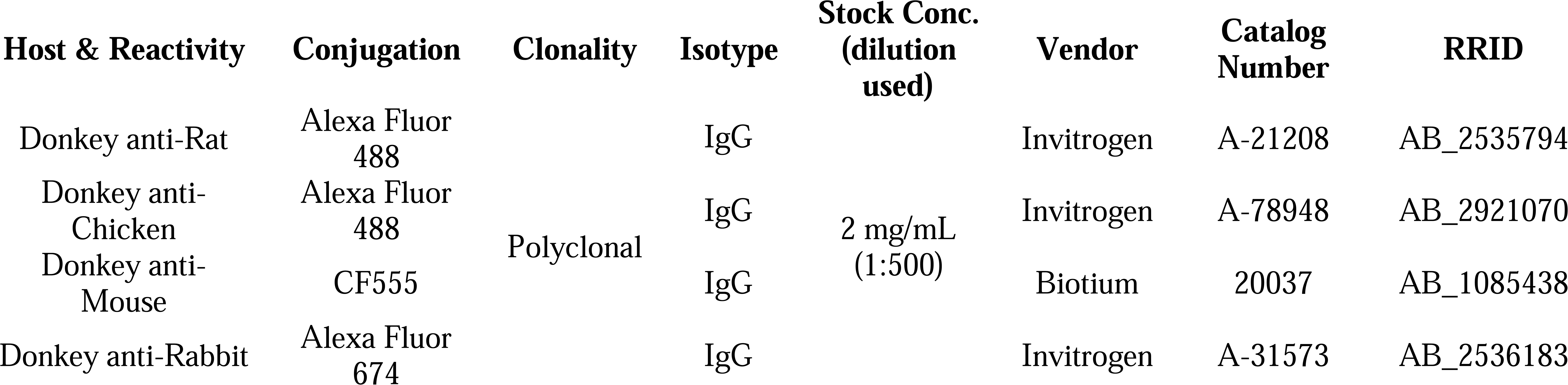
Summary of secondary antibodies used in the study. Listed are the host species and reactivity, conjugation, clonality, stock concentrations and working dilutions, as well as the vendor and catalog numbers for each secondary antibody.

**Table S3.**
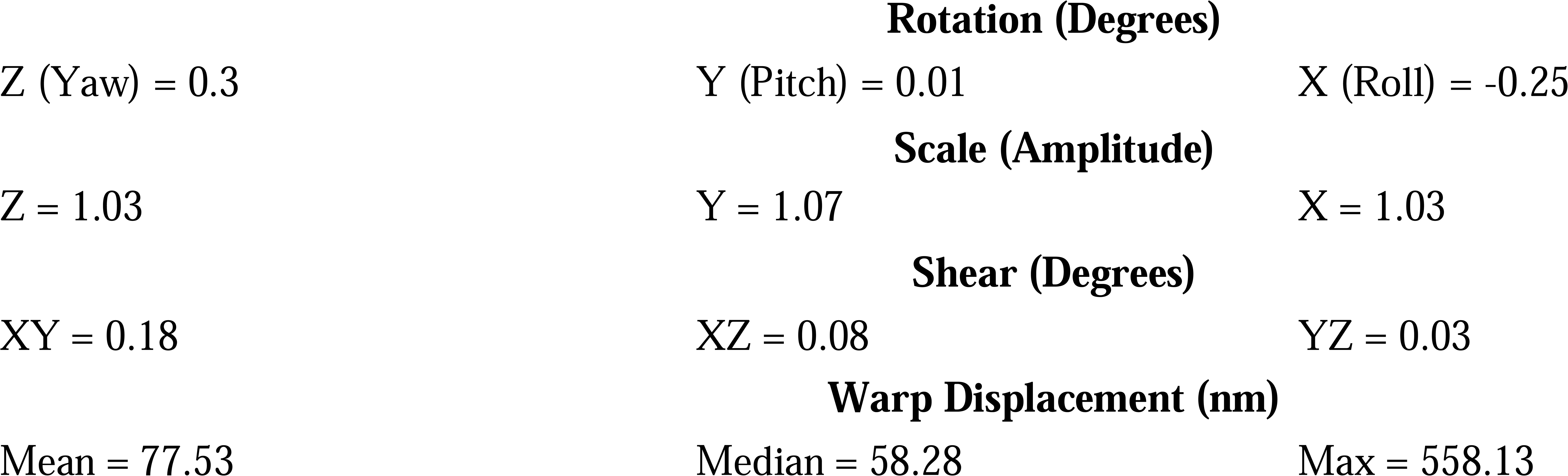
Affine registration metrics. Reported values include shear, rotation, and scale parameters, as well as the mean, median, and maximum displacement measured for the non-linear warp transform. Data were acquired with the oblique plane microscope.

**Table S4.**
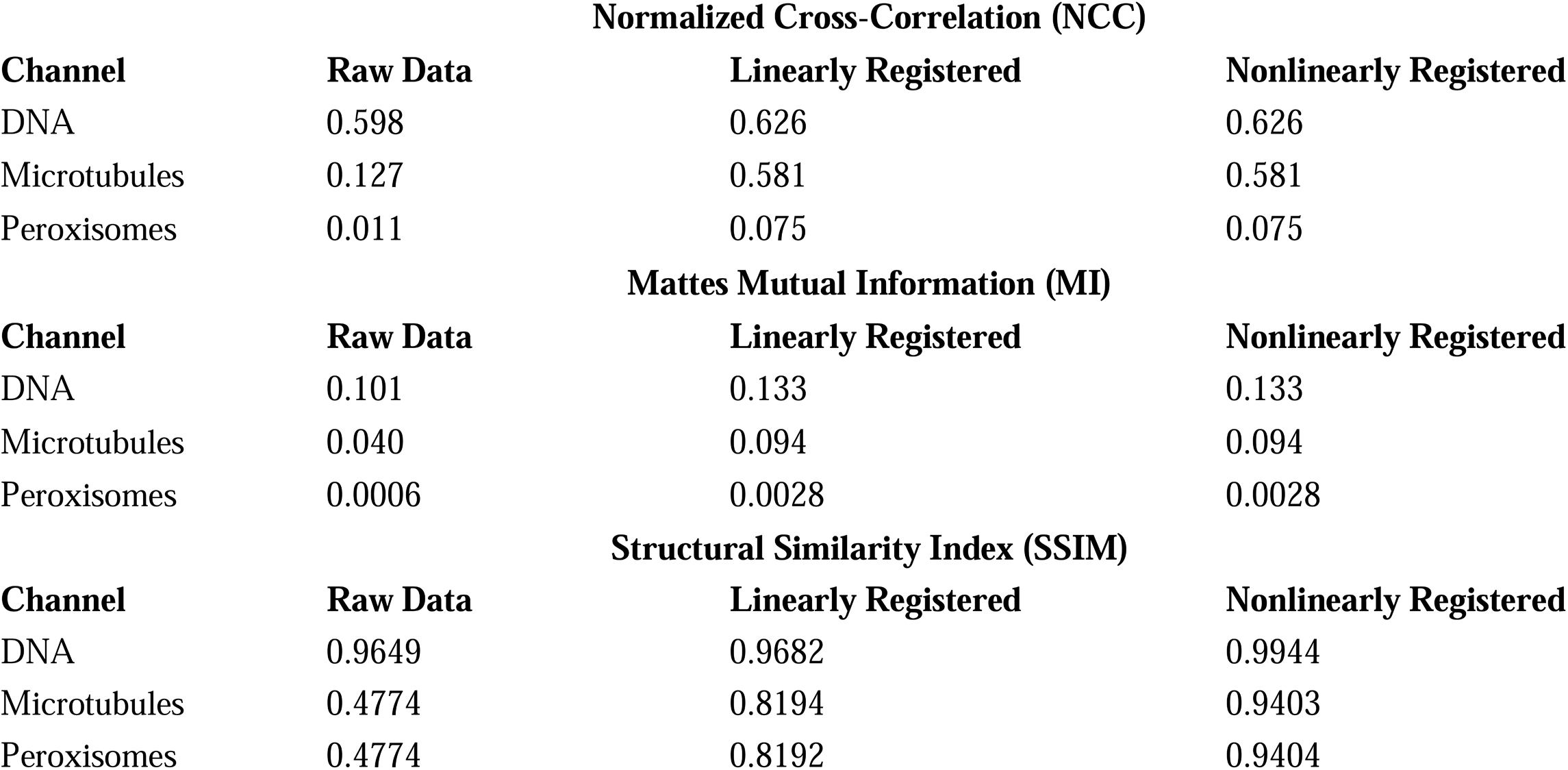
Quantitative evaluation of registration accuracy using intensity– and structure-based similarity metrics. Normalized cross-correlation (NCC) and Mattes mutual information (MI) were computed using the ANTsPy package with regular sampling and a 50% voxel sampling fraction. Structural similarity index (SSIM) was computed using the scikit-image package. Metrics are reported for raw (unregistered) data, after linear (affine) registration, and after additional non-linear registration for each imaging channel. While linear registration substantially improved global similarity metrics, non-linear registration did not further increase these global similarity metrics. SSIM likewise showed no improvement with non-linear warping and, in this dataset, decreased after registration, consistent with SSIM’s sensitivity to intensity scaling, background differences, and local contrast changes introduced by interpolation and resampling. Together, these results suggest the interpretation that non-linear registration primarily corrects localized geometric distortions that may not be captured by global intensity-based similarity measures.

**Table S5.**
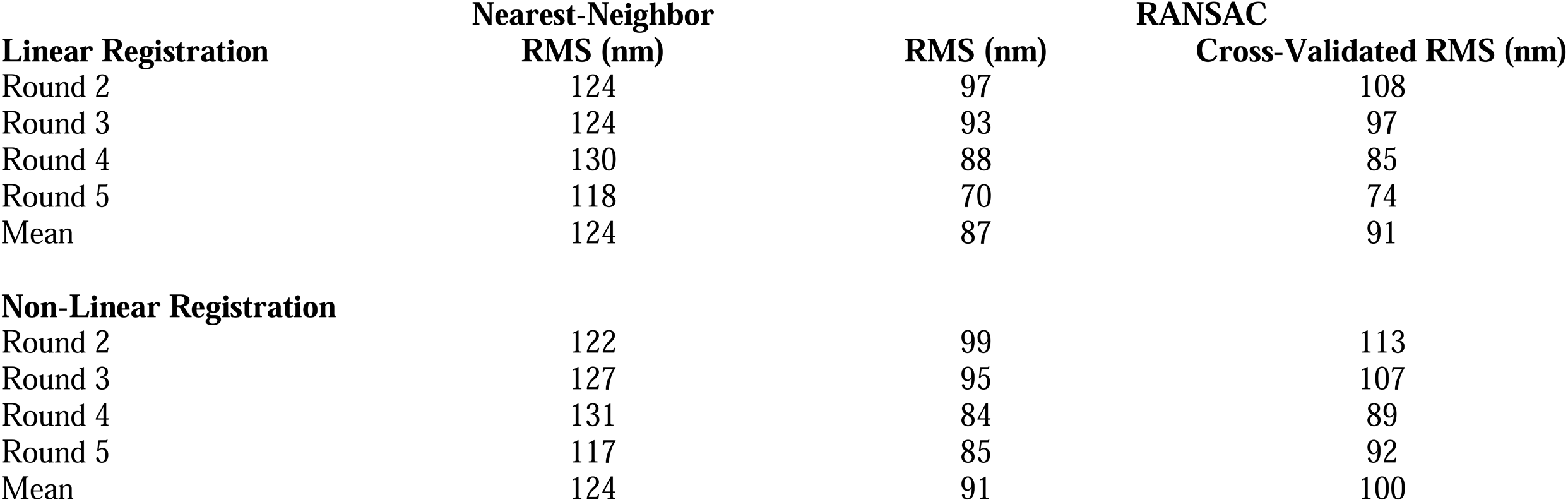
Independent estimates of registration error using mitochondrial landmarks across iterative imaging rounds. Registration accuracy was quantified in the mitochondrial channel after microtubule-guided registration. For each moving round (2–5) relative to round 1, a target-registration-error–like proxy was computed using (i) mutual nearest-neighbor pairing of detected mitochondrial puncta and reporting the RMS paired distance, and (ii) a descriptor-based RANSAC approach that fits a global similarity transform to inlier matches, reporting both the inlier RMS residual and a cross-validated RMS residual on held-out inliers. Metrics are shown for linear registration alone and for combined linear plus non-linear registration. Distances are reported in nanometers.

**Table S6.**
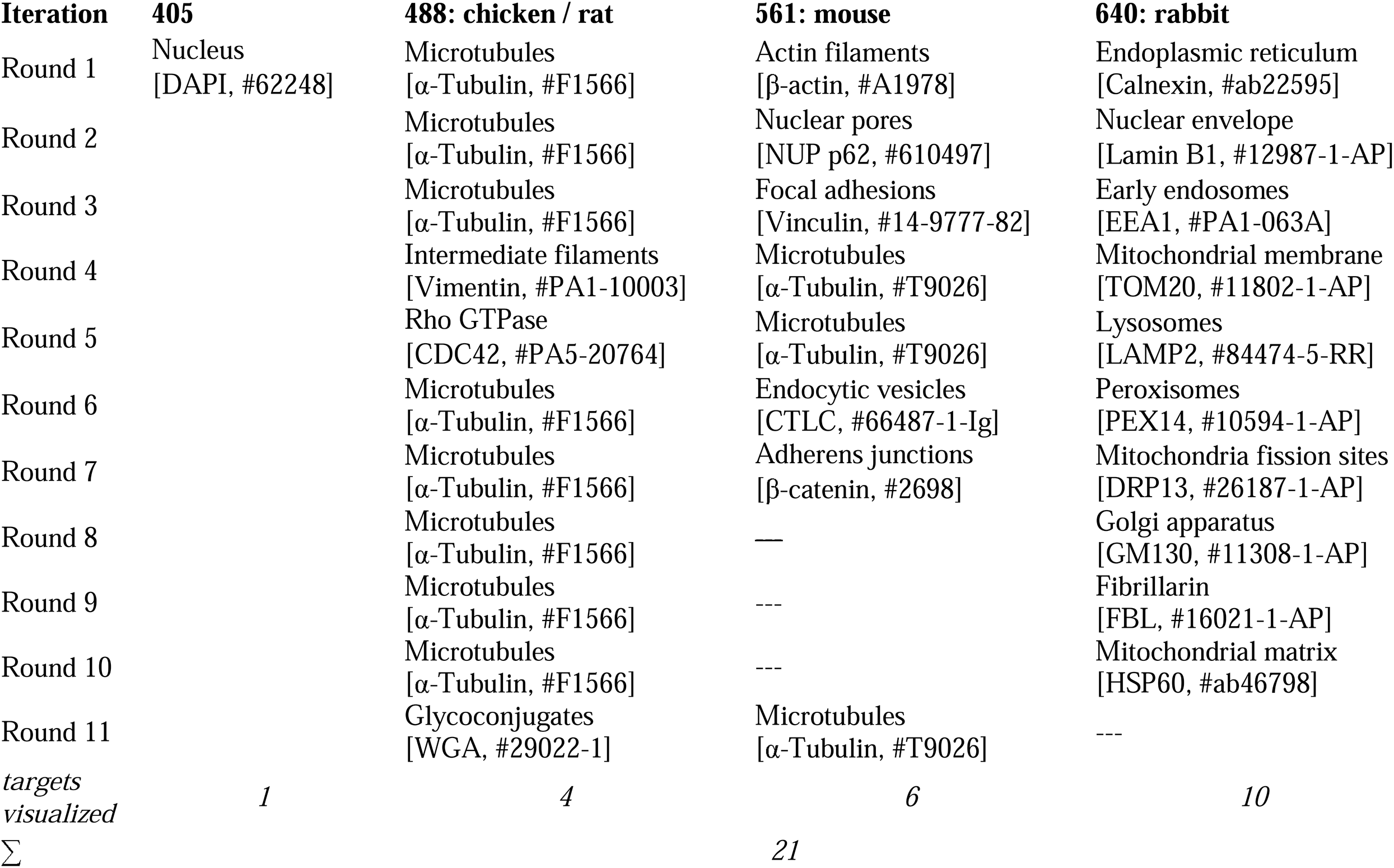
Multiplexed labelling experimental design for visualization of subcellular organization. Experimental design showing the antibody host species, fluorophore excitation channels (405, 488, 555, and 647 nm), and corresponding targets imaged across eleven labeling rounds. The 488 nm or 555 nm channel was used as a registration channel throughout all cycles. At the bottom, the number of targets visualized per channel is summarized, along with the total number of distinct labels imaged across the full multiplexing experiment.

**Table S7.**
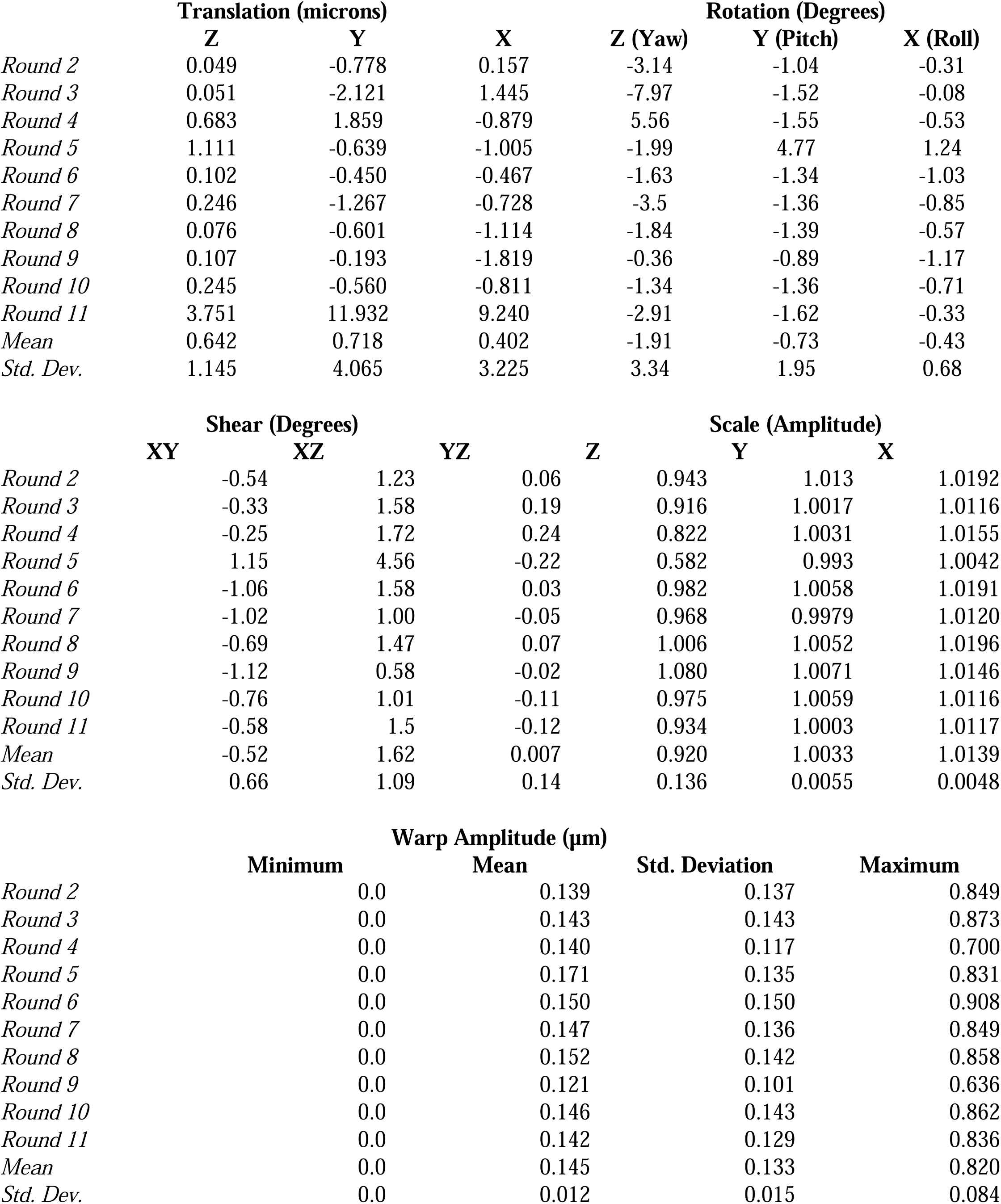
Quantitative registration parameters across iterative imaging rounds. Reported values summarize the decomposition of affine and non-linear registration transforms between round 1 and subsequent imaging rounds. Parameters include translation (µm), rotation (degrees), scale (unitless), and shear (degrees) derived from the affine transform, as well as the mean, maximum, and standard deviation of the displacement field from the non-linear warp. Mean and standard deviation values across rounds are provided to quantify the magnitude and variability of round-to-round geometric transformations.

**Table S8.**
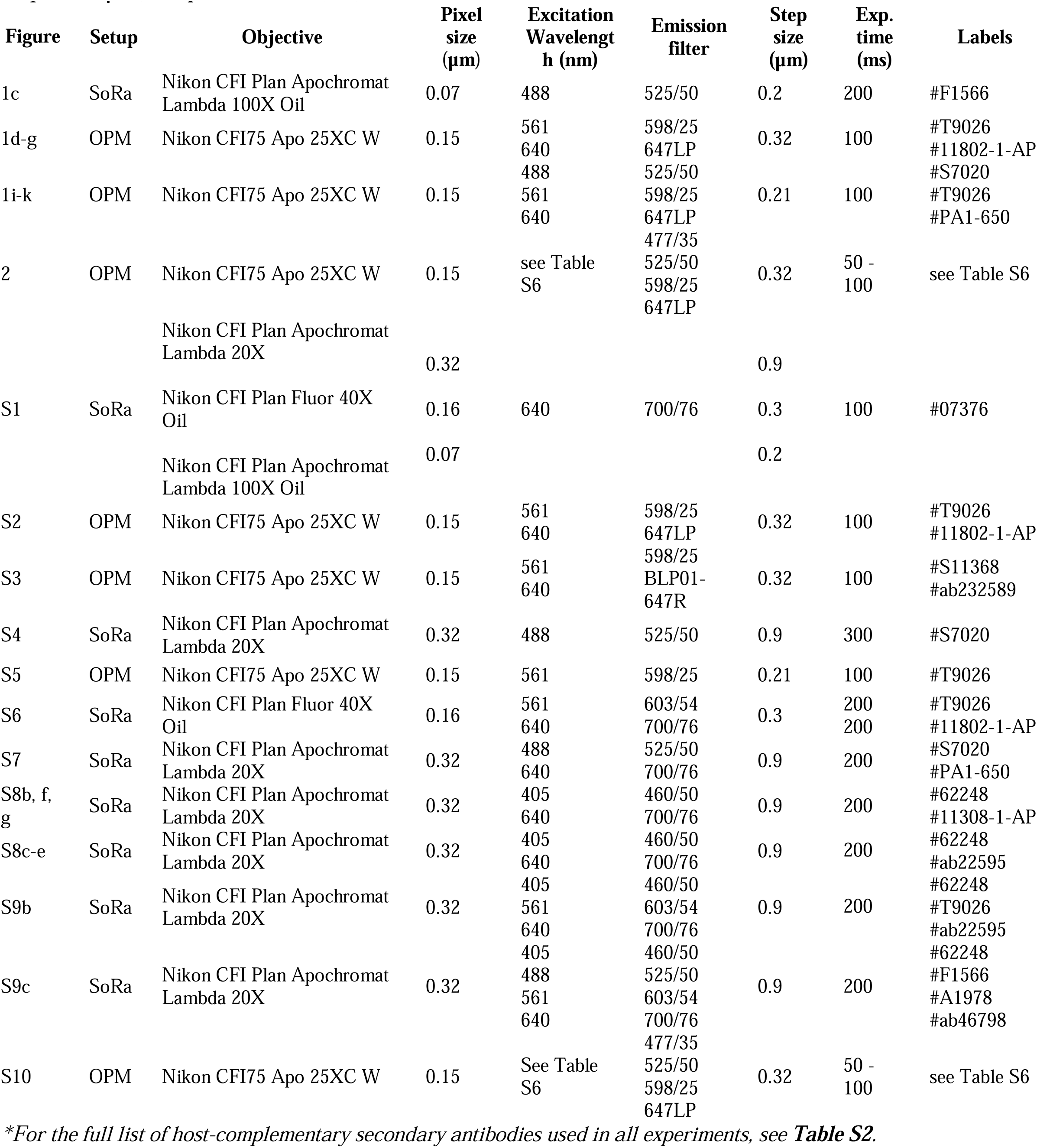
Imaging Parameters. Detailed summary of imaging conditions used on the pinning disk confocal microscope (SoRa) and oblique plane microscope (OPM), including objective specifications (immersion type and magnification), physical pixel size (µm), wavelengths (nm), emission filter type, z-step size (µm), exposure time (ms), and labels used.

## SUPPLEMENTARY NOTES

### Supplementary Note 1 – Strategies for Multiplexed Immunostaining in Expansion Microscopy

Expansion Microscopy (ExM) has rapidly evolved since its first description, providing a powerful means to achieve nanoscale imaging with conventional microscopes. The original protocol, however, relied on pre-expansion labeling combined with harsh proteolytic digestion, which destroyed most proteins and left only covalently anchored fluorophores as “phantoms” of the original structures^24^. While effective for visualization of a limited number of targets (typically on the order of a few), this approach was inherently incompatible with highly multiplexed labeling strategies. To address these limitations, numerous ExM variants have been developed with alternative gel chemistries, denaturation strategies, and labeling schemes, greatly broadening its utility for imaging single cells, tissue sections, and even entire organisms.

Today, immunostaining in ExM generally falls into two camps: pre-expansion labeling, in which targets are stained before gelation and expansion, and post-expansion labeling, where staining is performed after the specimen has been expanded. To perform multiplexing in a format compatible with pre-expansion immunostaining, the sample must be stained simultaneously with all primary antibodies prior to embedding. Because the number of orthogonal antibody pairs is limited (e.g., goat, mouse, rabbit, etc.), alternative readout mechanisms are required. The most common strategy is to use oligo-conjugated antibodies, where the attached DNA sequence can be decoded in a cyclic format after expansion^25^. Often, antibody conjugation must be performed independently, increasing experimental complexity. While this enables multiplexing, direct immunofluorescence provides only limited signal amplification, which is is particularly important in ExM because fluorophore density is defined per unit volume; since the number of labeled molecules is fixed while the specimen volume increases by the cube of the linear expansion factor (E^3^), the effective fluorophore concentration decreases proportionally (∝1/E^3^). Some improvement can be achieved by increasing labeling stoichiometry per target (e.g., using pre-complexed primary antibodies with labeled secondary reagents such as F(ab’)_2_ fragments or nanobodies) or by adopting branched-DNA hybridization schemes to boost readout signal. And, if the oligo is further modified to include a primary amine, it can be covalently incorporated into the hydrogel during polymerization^24^, ensuring retention of the readout sequence throughout expansion. To prevent the hydrogel from shrinking during secondary oligo hybridization, which requires salt-containing buffers, samples are in some implementations embedded in a secondary, non-expandable hydrogel; however, this introduces diffusion barriers for sequential labeling and alters the refractive index, exacerbating spherical aberrations.

In contrast, post-expansion labeling offers several advantages for multiplexed imaging. Because staining is performed after gelation, trapped or nonspecifically bound fluorophores can be washed away, reducing background. The porous nature of the expanded gel also facilitates deep and uniform antibody penetration^26^, even for large complexes such as IgG (∼150 kDa, ∼14.5 nm), a feature particularly beneficial for labeling tissues and organs where diffusion barriers are significant and labeling time is greatly reduced. Importantly, indirect pre-expansion immunofluorescence (primary + secondary antibodies) introduces a linkage error of ∼15–20 nm in the unexpanded state, which scales proportionally with the expansion factor; post-expansion labeling minimizes this error and is further compatible with smaller affinity reagents such as Fab fragments or nanobodies, making it especially valuable when performing iterative ExM. A key consideration for post-expansion labeling is the preservation of antigenicity, as the gelation and denaturation steps can disrupt epitopes and reduce labeling efficiency. The primary methods for integrating proteins into the hydrogel are through covalent anchoring with AcX or polymer entanglement. However, covalent anchoring can mask epitopes and reduce labeling efficiency, whereas polymer entanglement, used here, retains proteins while better preserving epitope accessibility.

### Supplementary Note 2 – Considerations for Fixation and Mechanical Relaxation in ExM

Expansion microscopy workflows, whether relying on pre– or post-expansion labeling, share critical dependencies on how specimens are fixed and mechanically relaxed, as these steps directly govern protein retention, antigen accessibility, and ultimately the quality of downstream labeling. Fixation with different ratios of paraformaldehyde (PFA) and acrylamide (AA) represents a trade-off between epitope preservation, protein retention, and fluorescence stability: higher PFA increases crosslinking and protein retention but often masks epitopes, whereas lower PFA or delayed AA introduction improves antigen accessibility, as demonstrated in eMAP^13^ and U-ExM^27^. Similarly, the step used to relax tissue mechanics, whether proteolysis with proteinase K or heat/SDS denaturation, has major consequences for labeling. Proteolysis efficiently softens tissues but removes many proteins, limiting downstream detection, while heat-based denaturation preserves more protein content, albeit with a risk of partial epitope disruption. Thus, regardless of whether labeling occurs before or after expansion, careful balancing of fixation chemistry and relaxation method is critical for maximizing both retention and accessibility of antigens. To preserve the maximum number of proteins, we applied an U-ExM^14,27^-inspired workflow, which relies on mild heat denaturation of the specimen and allows for detection of antigens in the post-expansion state.

### Supplementary Note 3 – Quantitative Evaluation of Registration Accuracy

To assess the fidelity of image alignment across iterative labeling cycles, we employ a suite of complementary registration metrics that capture both geometric and intensity-based correspondence between volumes. These methods are briefly summarized below.

#### Target Registration Error (TRE)

TRE is a landmark-based metric that directly measures geometric misalignment. It is defined as the 3D Euclidean distance between corresponding image features after applying the registration transform. Specifically, given N landmark pairs, each with coordinates (*x_i_*, *y_i_*, *z_i_*) in the reference image and corresponding coordinates 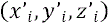 in the moving (transformed) image, the TRE for each pair is calculated as:

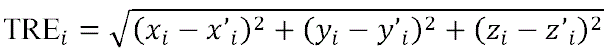

TRE reflects how well independent structures align following registration and is typically summarized using mean, median, and range. This provides a direct and interpretable measure of how precisely structures are aligned across rounds. In this work, because landmark correspondence is inferred from puncta matching rather than a known a priori, the reported values represent a TRE-like proxy rather than a ground-truth TRE.

#### Normalized Cross-Correlation (NCC)

NCC quantifies the linear correlation between voxel intensities in the two volumes and is calculated as the Pearson correlation coefficient across all voxel pairs. For two images *I*_1_ and *I*_2_ with N voxels each:

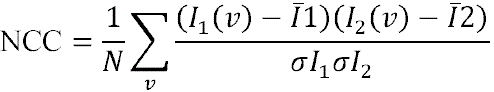

NCC ranges from –1 to 1, where 1 indicates perfect alignment of structural intensities and 0 indicates no correlation. In high-resolution fluorescence microscopy of the same target structure, well-registered volumes typically achieve NCC values above 0.9. Because it assumes a linear relationship between intensity values, NCC underperforms when fluctuations in the expansion coefficient or staining efficiency result in intensity differences between the two images (see **Figure S6**).

#### Mutual Information (MI)

MI is an information-theoretic measure of how much knowing the intensity in one image reduces uncertainty about the other. Unlike NCC, MI does not assume a linear relationship and is well-suited for variably scaled data, such as those arising from differences in the expansion coefficient between imaging rounds. Like NCC, MI remains sensitive to fluctuations in staining efficiency^28^. It is calculated from the joint intensity histogram of the two volumes:

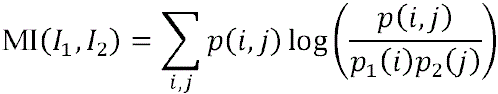

where *p*(*i*, *j*) is the joint probability of observing intensity in image *I*_1_ and *j*. in *I*_2_, and *p*_1_ and *p*_2_ are the marginal distributions. MI increases when structural alignment leads to a more peaked joint histogram and decreases when images are misaligned. While absolute MI values are less interpretable, improvements in MI after registration indicate enhanced alignment.

#### Structural Similarity Index (SSIM)

SSIM evaluates perceptual similarity between images by comparing local patterns of luminance, contrast, and structural texture. For local windows centered at each voxel:

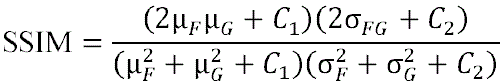

where µ, σ, and σ*_FG_* are local means, standard deviations, and cross-covariance terms, respectively, and *C*_1_, *C*_2_ are stability constants. SSIM results range from –1 to 1: a value of 1 indicates perfect structural similarity, while –1 represents perfect anti-correlation (e.g., an image and its negative). SSIM is particularly useful for assessing registration accuracy in high-resolution microscopy data, where preserving local spatial relationships is critical. Using the default implementation in Scikit-image, we compute SSIM over 3D neighborhoods to capture structural alignment in volumetric datasets.

### Supplementary Note 4 – Limitations and Future Directions

#### Limitations

Despite its advantages, Cy-ExM has several limitations that warrant consideration. First, antibody compatibility remains a bottleneck, and certain epitopes may be sensitive to cryo-fixation or expansion-associated processing. Because our workflow relies on indirect immunofluorescence, each cycle requires orthogonal primary–secondary antibody pairs, which constrains panel design and multiplexing depth per round. This limitation could be alleviated by adopting direct labeling strategies (e.g., directly conjugated primaries) or using pre-complexed Fab–antibody assemblies to expand the number of usable channels without increasing cross-reactivity. Second, although Cy-ExM preserves antigenicity for many targets, fluorescent proteins are variably preserved during cryo-fixation and denaturation; when intrinsic fluorescence is critical, additional optimization or post-expansion immunostaining is likely required. Third, while β-mercaptoethanol provides a relatively gentle mechanism for antibody removal, repeated reduction cycles may still affect the stability of particularly sensitive epitopes or labeling chemistries, and the practical ceiling on cycle number may depend on target class and specimen type. Finally, nanoscale 3D imaging across multiple cycles generates large volumetric datasets, necessitating substantial storage and computational resources as well as automated workflows for registration, segmentation, and downstream analysis. Addressing these limitations through expanded labeling chemistries, continued optimization of fixation/denaturation/stripping conditions, and more efficient computational pipelines will further broaden the applicability and scalability of Cy-ExM.

#### Future directions

Looking ahead, Cy-ExM establishes a foundation for several promising extensions. Combining our cyclic workflow with iterative expansion microscopy could push effective resolution into the ∼20 nm regime, enabling more detailed mapping of protein nanodomains, cytoskeletal architectures, and organelle interfaces. Future implementations could also substantially increase the molecular information captured per cycle by integrating automated fluidics, broadband illumination sources, tunable emission filters, ultra-narrowband fluorophores, and improved spectral unmixing algorithms. Together, these advances could move Cy-ExM toward substantially higher multiplexing depth (e.g., ∼80-plex and potentially >100-plex, depending on channel allocation, fluorophore performance, and cycle count) while maintaining image fidelity. In addition, the approach should be adaptable to more complex biological specimens, including organoids and tissue sections, particularly when paired with cryo-preservation strategies that maintain subcellular detail at depth. Ultimately, the ability to visualize molecular organization across dozens of targets within intact 3D specimens holds promise for systematically characterizing how subcellular protein architecture varies across cell types and states, and how compartment-level organization relates to biological function across cellular, organelle, and tissue scales.

